# Cross-Study Transcriptomic Meta-Analysis Reveals Conserved Adaptive Programs in Escherichia coli K-12

**DOI:** 10.64898/2026.08.20.745921

**Authors:** Mohammad Javad Golmohammadi

## Abstract

Adaptive laboratory evolution (ALE) provides a powerful framework for investigating the molecular basis of bacterial adaptation, yet the extent to which transcriptional responses recur across independent evolutionary trajectories remains poorly understood. Here, we performed a cross-study transcriptomic meta-analysis of *Escherichia coli* K-12 ALE experiments conducted under diverse genetic and environmental selective conditions. Seven study-level inputs were integrated, including a combined signature derived from three related *menF*-associated comparisons and six independent transcriptomic datasets. Study-specific transcriptional responses were harmonized according to their direction and statistical evidence, followed by rank-based meta-analysis to identify genes showing recurrent expression changes across evolutionary contexts. We identified 109 conserved core genes, comprising 32 upregulated and 77 downregulated genes, that were supported across the majority of independent study-level inputs. Functional enrichment and protein-protein interaction analyses revealed that these conserved responses were organized into distinct biological modules, with prominent representation of flagellar assembly, chemotaxis, and motility, together with transport and curli/biofilm-associated functions. Highly connected genes included *fliC, fliA, cheA, cheB, cheW, cheY, motA,* and *motB* within the flagellar and chemotaxis-associated network, and *csgA, csgD, csgE, csgF,* and *csgG* within the curli-associated module. Overall, these findings demonstrate that, despite substantial diversity in evolutionary conditions and trajectories, *E. coli* adaptation is accompanied by a reproducible transcriptional component involving coordinated remodeling of motility, environmental sensing, transport, and surface-associated functions. Cross-study integration of ALE transcriptomes therefore provides a framework for distinguishing recurrent features of bacterial adaptation from context-specific transcriptional responses.

Graphical Abstract
Overview of the analytical strategy used to identify common transcriptional features of adaptive evolution in *Escherichia coli* K-12.
Transcriptomic profiles from independent ALE experiments were harmonized and integrated to identify genes showing recurrent directional changes across evolutionary conditions. The resulting conserved gene set was further examined using functional enrichment and protein-interaction analyses, followed by hub-gene and network-module characterization. These complementary analyses highlighted recurrent changes in motility and chemotaxis, nutrient acquisition, and surface-associated functions across diverse evolutionary trajectories.

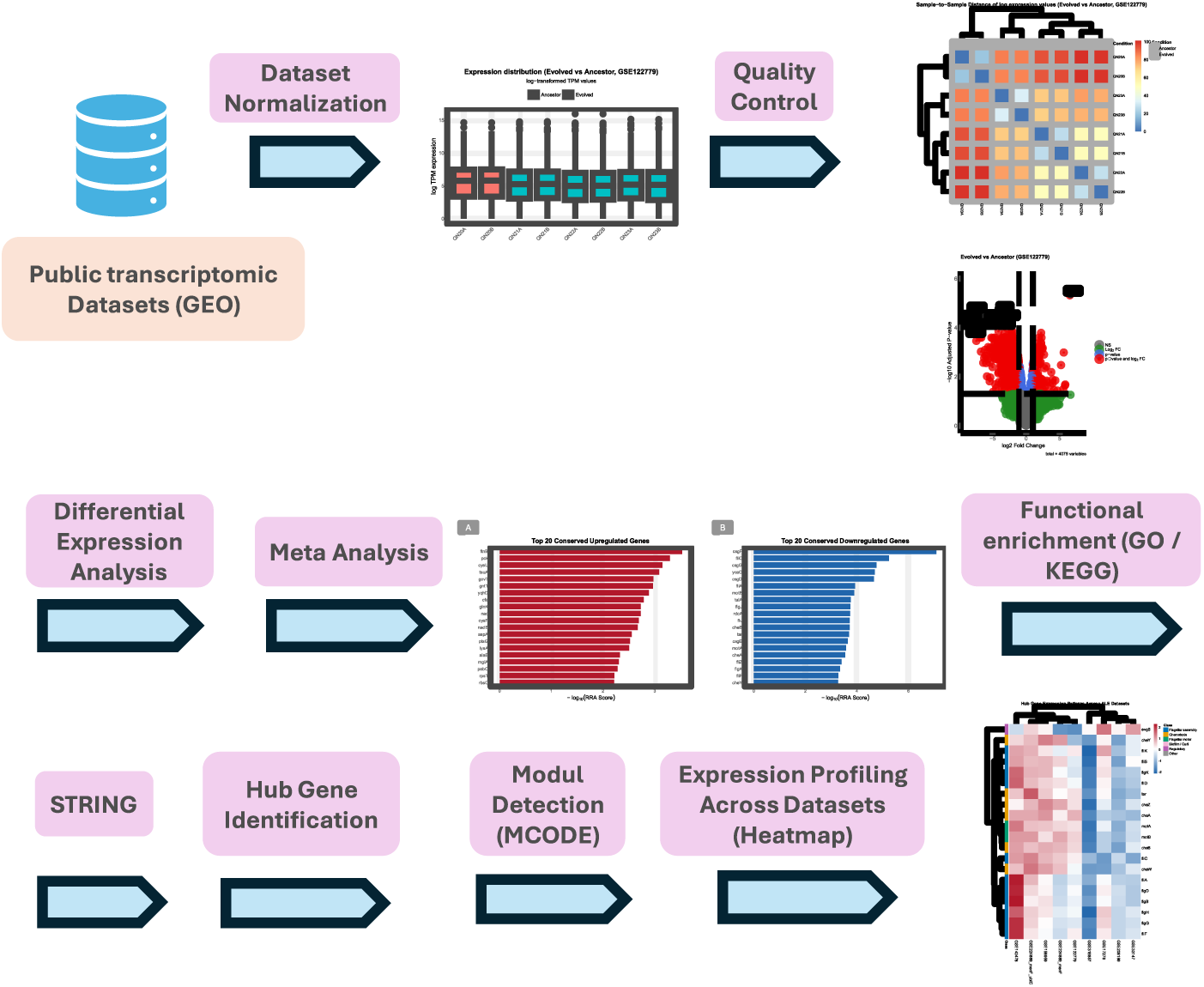

## 1. Introduction

Microbial populations can rapidly adapt to environmental and genetic perturbations through the accumulation of heritable changes that increase fitness under selection. Because bacteria have short generation times and can be propagated for many generations under controlled conditions, experimental evolution provides a tractable framework for studying adaptation over time (Elena & Lenski, 2003; Kawecki et al., 2012) Adaptive laboratory evolution (ALE) enables the application of defined selective pressures to microbial populations and the subsequent characterization of genetic, physiological, and molecular changes associated with adaptation (Dragosits & Mattanovich, 2013) *Escherichia coli* is particularly well suited to ALE because of its extensively characterized genome, powerful genetic and molecular toolkit, rapid growth, and the availability of numerous independently evolved populations and long-term experimental evolution datasets (Cronan, 2014; Kurokawa & Ying, 2019; Lenski, 2017).

A central insight emerging from experimental evolution is that adaptation can involve both highly reproducible and strongly context-dependent responses (Cooper et al., 2003; Fong et al., 2005). Parallel populations exposed to the same selective environment can converge on similar phenotypes despite differences in their underlying genetic and transcriptional states (Favate et al., 2022; Fong et al., 2005). Early studies of parallel *E. coli* evolution demonstrated that independently evolved populations can reach convergent growth phenotypes while retaining substantially different global gene-expression states, with only a subset of expression changes consistently shared among populations (Fong et al., 2005). These observations suggest that the transcriptional landscape of adaptation contains at least two components: a broad context-dependent component associated with compensatory or lineage-specific remodeling, and a smaller conserved component that may represent recurrent solutions to common physiological constraints (Cooper et al., 2003; Fong et al., 2005).

The distinction between conserved and context-specific responses is particularly important because ALE has now been applied to a wide range of evolutionary challenges (Sandberg et al., 2019). *E. coli* populations have been experimentally adapted to altered carbon sources, metabolic deficiencies, chemical stresses, elevated temperatures, and other environmental or genetic perturbations. Such experiments have demonstrated that adaptation can involve extensive metabolic rewiring, changes in transcriptional regulation, altered transport capacity, modification of stress-response systems, and remodeling of cellular resource allocation (Dragosits & Mattanovich, 2013; Franchini & Egli, 2006; McCloskey et al., 2018; Rychel et al., 2025; Sandberg et al., 2019). For example, ALE of genome-reduced *E. coli* revealed extensive metabolic rewiring and transcriptome-wide remodeling associated with restoration of growth, with changes in global transcriptional regulation contributing to the reorganization of cellular metabolism (Choe et al., 2019).

Likewise, recent work on thermal adaptation illustrates the systems-level complexity of transcriptional evolution. Independently evolved *E. coli* strains capable of growth at temperatures that are lethal to the ancestral strain exhibited coordinated changes involving stress responses, metabolism, iron acquisition, fimbrial functions, and flagellar systems. Importantly, this study showed that transcriptomic analysis can reveal coherent adaptive mechanisms despite the complexity of the underlying genetic changes (Rychel et al., 2025). These findings reinforce the concept that evolutionary adaptation should not be interpreted solely as a collection of individual mutations or differentially expressed genes; rather, adaptive phenotypes can emerge through coordinated remodeling of interconnected cellular systems.

Despite the growing number of *E. coli* ALE transcriptomic studies (Peng et al., 2025), however, transcriptomic analyses have generally been conducted within individual evolutionary experiments and their specific selective contexts. Each experiment is generally designed around a particular selective pressure, genetic background, or biological hypothesis, and differential expression is consequently evaluated within that experimental context (Hirasawa & Maeda, 2022; Puentes-Téllez et al., 2014; G. Wang et al., 2023). This approach is appropriate for identifying mechanisms specific to an individual evolutionary trajectory, but it makes it difficult to determine whether an observed transcriptional response represents a general feature of adaptation or simply a consequence of the particular environment in which evolution occurred (Puentes-Téllez et al., 2014). Differences in experimental design, sequencing or microarray platform, normalization procedures, sample sizes, evolved lineages, and statistical analysis further complicate direct comparison among studies (Chen et al., 2011; Olivas-Bernal et al., 2026).

A cross-study meta-analysis provides an opportunity to address this limitation by shifting the analytical question from “which genes change in this experiment?” to “which transcriptional changes recur across independent evolutionary experiments?” (Olivas-Bernal et al., 2026) Such an approach is particularly valuable for ALE because evolutionary trajectories are inherently variable. A gene that changes strongly in one evolved population may not be altered in another population even when both populations achieve similar phenotypic outcomes. Conversely, genes or pathways that repeatedly change across genetically and environmentally distinct evolutionary experiments may represent more fundamental components of bacterial adaptation (Kavvas et al., 2022; Kram et al., 2017). Identifying these recurrent responses can therefore help distinguish general adaptive programs from selection-specific transcriptional signatures (Kavvas et al., 2022; Olivas-Bernal et al., 2026).

Although individual ALE studies have provided valuable insights into the molecular basis of adaptation, the extent to which specific transcriptional changes are reproducible across distinct evolutionary trajectories remains largely unresolved. The large number of independently generated datasets provides an opportunity to address this question systematically, but their study-specific nature makes it difficult to distinguish recurrent adaptive responses from changes associated with particular selection regimes (Dalldorf, Hefner, et al., 2024; Kim et al., 2024; Zion et al., 2024). Identifying genes that repeatedly exhibit concordant transcriptional changes across independent evolutionary experiments could therefore reveal a conserved component of the adaptive response that is not readily apparent from individual studies (Rychel et al., 2025).

Such a conserved transcriptional signature would provide a complementary perspective on bacterial adaptation by identifying genes whose altered expression is reproducible despite differences in the selective environment and evolutionary history (Kavvas et al., 2022). Importantly, recurrent gene-level responses can subsequently be examined at the functional and network levels to determine whether they represent coordinated biological processes (Dalldorf, Hefner, et al., 2024; Rychel et al., 2025; Saxena et al., 2025). This hierarchical approach—from conserved genes to functional enrichment and interaction networks—can link the reproducibility of individual transcriptional responses to their broader biological significance (Kavvas et al., 2022; Li et al., 2026; Rychel et al., 2025; Saxena et al., 2025).

Taken together, these considerations motivate a systematic cross-study investigation of transcriptional adaptation in *E. coli* K-12. The availability of independent ALE transcriptomic datasets generated under diverse evolutionary conditions provides an opportunity to identify transcriptional responses that are reproducible beyond the boundaries of individual experiments (Abdel-Salam et al., 2023; Özel et al., 2024). In the present study, we therefore integrated publicly available transcriptomic datasets representing distinct evolutionary contexts and evaluated the consistency of gene-level responses across these independent trajectories. We hypothesized that, despite substantial variation among experimental conditions and evolutionary histories, a subset of genes and functional systems would exhibit recurrent directional changes indicative of conserved adaptive remodeling (Rychel et al., 2025).

Our primary objective was to identify and characterize this conserved transcriptional component of laboratory adaptation in *E. coli* K-12. We further sought to determine whether recurrently altered genes are organized into biologically coherent pathways and interaction networks, thereby providing insight into the cellular systems that are repeatedly remodeled during adaptation. By integrating evidence across independent ALE experiments rather than interpreting each evolutionary trajectory in isolation, this study aims to distinguish broadly recurrent adaptive responses from context-dependent transcriptional changes and to provide a systems-level view of the transcriptional architecture underlying bacterial adaptation.

## 2. Experimental Procedures

### 2.1. Study design and transcriptomic datasets

A cross-study transcriptomic meta-analysis was performed to identify genes exhibiting recurrent and directionally consistent expression changes across independent adaptive laboratory evolution (ALE) experiments in *Escherichia coli* K-12. Publicly available transcriptomic datasets were selected on the basis of three principal criteria: (i) the study involved an evolved *E. coli* population or strain generated through laboratory adaptation; (ii) transcriptomic measurements were available for comparison between an evolved population and its corresponding ancestral, parental, or non-evolved reference state; and (iii) sufficient gene-level expression information was available to derive a study-specific directional transcriptional signature.

The final meta-analysis incorporated seven study-level inputs. Three related comparisons associated with *menF*-deficient evolution were first integrated into a single combined signature and subsequently treated as one independent input in the cross-study meta-analysis. The remaining six inputs were derived from GSE140478 (Rychel et al., 2025), GSE158959 (Babel & Krömer, 2020), GSE17276 (Kinnersley et al., 2009), GSE206196 (Alkim et al., 2022), GSE33147 (Fong et al., 2005), and GSE316857 (Patil et al., 2026). These datasets represented diverse evolutionary contexts, including adaptation following metabolic gene perturbation, isoprenol-associated selection, long-term glucose limitation, glycerol adaptation, homoserine-associated evolution, and paraquat-associated evolution. The thermal-adaptation dataset GSE140478 was restricted to the 44°C condition to maintain a single, biologically coherent evolutionary comparison. Likewise, only the glycerol-evolution arm of GSE33147 was included. For GSE17276, consortium samples were excluded and only the monoculture evolution experiments were retained.

The three *menF*-associated comparisons comprised the *menF* knockout versus its independently adapted derivatives in GSE224889, the *menF/ubiC* knockout versus adapted derivatives in GSE224889, and the evolved Δ*menF*Δ*entC* populations in GSE122779 (Anand et al., 2019). These three related signatures were integrated before the principal meta-analysis to avoid treating closely related comparisons as three independent study-level observations. The resulting combined signature was subsequently considered a single meta-analysis input. The final RRA analysis comprised seven study-level inputs (Signed Z-score files): one combined *menF*-associated signature derived from three related comparisons, and six independent transcriptomic signatures derived from GSE140478, GSE158959, GSE17276, GSE206196, GSE33147, and GSE316857.

### 2.2. Quality control and visualization

Quality control analyses were performed using the normalized or processed expression matrices to assess sample-level consistency and identify potential technical abnormalities. Boxplots were generated to examine the distribution of gene expression values across samples and to verify that the samples showed comparable overall expression distributions. Consistent and appropriately aligned boxplot distributions were considered indicative of satisfactory normalization and absence of substantial global expression bias among samples.

Principal component analysis (PCA) was performed to evaluate the major sources of variation among samples and to determine whether biological groups exhibited coherent transcriptional separation. PCA plots were inspected for clustering of ancestral and evolved samples and for the presence of samples showing substantial deviation from their respective biological groups. Hierarchical clustering and sample-level heatmaps were additionally used to assess the similarity structure among samples and to identify unexpected sample relationships.

Following differential-expression analysis, volcano plots were generated to visualize the relationship between estimated expression changes and statistical significance for each gene. These plots provided a global representation of the transcriptional response and facilitated identification of genes exhibiting both large effect sizes and strong statistical evidence. Differential-expression results were additionally summarized using MA plots, in which average expression abundance was plotted against estimated log fold change, allowing assessment of the distribution of expression changes across the range of gene abundance and visualization of potential intensity-dependent patterns.

All exploratory and differential-expression visualizations were generated using reproducible R-based workflows. The same general visualization framework was applied across datasets where appropriate, while the underlying transformation and statistical model were retained according to the characteristics of each dataset and its corresponding analytical pipeline.

### 2.3. RNA-sequencing data preprocessing and differential expression analysis

RNA-seq datasets were analyzed using DESeq2 when raw count matrices were available (Love et al., 2014) and using limma-based analysis when processed expression matrices were provided or when the dataset had already been represented in a normalized continuous-expression format (Law et al., 2014; Ritchie et al., 2015).

For GSE224889, the published count matrix was used to analyze the *menF* and *menF/ubiC* evolutionary comparisons separately. For the *menF* analysis, two *menF* knockout samples were compared with eight independently adapted samples, comprising four adapted populations with two biological replicates each. For the *menF/ubiC* analysis, the corresponding two knockout samples and eight adapted samples were selected from the same dataset. Gene-level count matrices were constructed from the published count table, duplicated gene identifiers were collapsed by summing counts across entries, and genes were retained when a count of at least 10 was observed in at least three samples. Differential expression was then evaluated using DESeq2 with a design formula of ∼ condition, with the ancestral knockout state specified as the reference level. Variance-stabilizing transformation (VST) was performed using blind = FALSE for sample-level quality assessment and visualization. GSE158959 was analyzed from the available gene-level expression/count data using DESeq2. The experimental design included three ancestral samples and nine evolved samples representing independently adapted *E. coli* populations associated with isoprenol adaptation. The condition factor was explicitly defined with the ancestral state as the reference level, and differential expression was estimated using a two-level design (∼condition). VST-normalized expression values were generated with blind = FALSE for downstream quality assessment. The resulting differential-expression table was retained with gene identifiers, log2 fold changes, and adjusted *P* values for subsequent meta-analysis.

For GSE122779, a processed log-transformed TPM expression matrix was obtained from the GEO-associated data file. The analysis was restricted to the Δ*menF*Δ*entC* evolutionary comparison, comprising two ancestral samples and six evolved biological replicates. Because the dataset was provided as a processed log-transformed expression matrix rather than raw sequencing counts, no additional count-based normalization was applied. Differential expression between evolved and ancestral samples was then assessed, and the resulting gene-level log fold changes and statistical significance values were converted to Signed Z-scores using the same procedure applied to the other study-level datasets. These Signed Z-scores were subsequently used to generate the ranked gene lists for cross-study Robust Rank Aggregation (Kolde et al., 2012).

For GSE140478, raw expression files were extracted and individually imported before being merged into a single gene-by-sample expression matrix. Gene identifiers containing numerical suffixes were standardized, and duplicated gene identifiers were collapsed by summing their expression values. Gene identifiers were subsequently mapped to *E. coli* K-12 MG1655 gene names using a reference GFF annotation, and duplicated gene symbols were again collapsed by summation. The analysis was restricted to samples corresponding to the 44°C evolutionary condition. This subset comprised two ancestral samples and 16 evolved samples representing eight independently thermally adapted strains. Lowly expressed genes were removed by retaining genes with expression values of at least 10 in a minimum of three samples (genes with zero or near-zero expression were excluded from all datasets prior to downstream analysis). Differential expression between ancestral and evolved samples was then assessed using DESeq2 with a two-level design (∼ condition), with the ancestral state specified as the reference level. Variance-stabilizing transformation (VST) was performed with blind = FALSE for downstream quality-control assessment and visualization.

### 2.4. Microarray preprocessing and differential expression analysis

Microarray datasets were processed according to the type and state of the available expression data. For GSE33147, raw Affymetrix CEL files were extracted and imported using the affy package (Gautier et al., 2004). Robust Multi-array Average (RMA) normalization was applied (Irizarry, 2003) to the complete CEL dataset, generating background-corrected, normalized, and log2-transformed expression estimates. Only the glycerol evolution condition was retained for the present analysis, corresponding to the comparison of evolved glycerol-adapted populations with their non-evolved reference state. Differential expression was subsequently assessed using limma (Ritchie et al., 2015).

GSE17276 was analyzed using the available processed log2 expression-ratio matrix, in which the reported values represented log2(Evolved/Ancestor) signals generated by competitive hybridization against a common ancestral reference. The analysis was restricted to monoculture evolution experiments, while consortium samples were excluded because they represented a distinct experimental context. The available monoculture measurements were retained, and genes were required to have measurements in at least 10 samples to ensure sufficient data coverage for downstream analysis. Gene identifiers were standardized and mapped to *E. coli* K-12 MG1655 gene annotations, with duplicated gene identifiers collapsed by averaging the available measurements. Because the input matrix already consisted of log2 expression ratios relative to a common ancestral reference, no additional between-array normalization was applied to the ratio values.

GSE206196 was analyzed using the normalized Agilent microarray expression matrix provided by the study. Six samples were retained for the comparison of the ancestral MG1655 wild-type state and the evolved 4E strain, with three biological replicates for each condition. The corresponding normalized expression matrix was analyzed using limma, with evolved 4E samples compared against the wild-type reference.

GSE316857 was analyzed using the published transcript-per-million (TPM) expression matrix, including three wild-type (WT) samples and three homoserine-adapted evolved (HPE) samples. Gene identifiers were standardized by removing numerical suffixes where applicable, and duplicate gene entries were collapsed by calculating the mean TPM value across corresponding entries. Gene identifiers were then mapped to *Escherichia coli* K-12 MG1655 gene annotations, with the original identifier retained when no corresponding gene name was available. Genes exhibiting TPM >1 in at least two samples were retained for downstream analysis. Because the expression matrix comprised continuous normalized expression measurements rather than raw sequencing read counts, differential expression analysis was performed using the limma linear modeling framework. The experimental design compared HPE with WT samples, with WT specified as the reference condition, and gene-wise linear models were fitted followed by empirical Bayes moderation of standard errors. Differential expression was quantified as the log2 fold change (HPE versus WT), and *P* values were adjusted for multiple testing using the Benjamini–Hochberg procedure (Benjamini & Hochberg, 1995). The resulting log2 fold changes and adjusted *P* values were subsequently used to calculate signed Z-scores, which were used to harmonize differential-expression effects across studies for the downstream cross-study meta-analysis.

### 2.5. Within-study differential-expression statistics and effect harmonization

For each dataset, gene-level differential-expression results were represented using the estimated log2 fold change (logFC) and multiple-testing-adjusted *P* value (adj.P.Val or the corresponding DESeq2 padj). For conventional DEG summaries, genes were considered significantly differentially expressed when the adjusted *P* value was below 0.05 and the absolute log2 fold change exceeded 1, where applicable. However, the cross-study meta-analysis was not based on binary DEG membership. Instead, the complete ranked transcriptional signal was retained to maximize information across heterogeneous datasets.

To harmonize statistical evidence across RNA-seq and microarray experiments, a Signed Z-score was calculated for each gene from its direction of change and adjusted significance. Adjusted *P* values below 1 × 10^-300 were truncated to 1 × 10^-300 to avoid numerical instability. The Signed Z-score was calculated as:

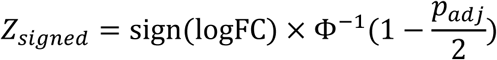

where (Φ^−1^) denotes the inverse cumulative distribution function of the standard normal distribution, (*p_adj_*) represents the Benjamini–Hochberg-adjusted *P* value, and (sign(logFC)) assigns the direction of the effect according to the corresponding log2 fold change. Thus, positive signed Z-scores indicated higher expression in the evolved condition relative to the corresponding reference, whereas negative values indicated lower expression.

Genes were subsequently ranked in descending order according to the absolute value of their signed Z-scores, thereby prioritizing genes with stronger statistical evidence for differential expression while retaining the direction of the effect. This transformation placed differential-expression evidence on a common statistical scale across datasets generated using different transcriptomic platforms and analytical pipelines, while preserving both the direction and relative strength of the evidence. Importantly, the subsequent meta-analysis was based on the ranks of the signed Z-scores rather than on direct comparisons of absolute expression measurements across platforms.

### 2.6. Integration of the three *menF*-associated comparisons

Because the three *menF*-associated comparisons represented related evolutionary contexts, their Signed Z-score profiles were integrated before inclusion in the main meta-analysis. Only genes present in all three input signatures were considered for this integration. The three Signed Z-scores were combined using a weighted Stouffer-type procedure (ZAYKIN, 2011). For the present analysis, the weight assigned to each comparison was defined as the square root of the total number of gene-level entries represented in its corresponding input file:

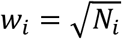

where (N_i_) denotes the total number of gene-level entries in input file. For each common gene, the combined score was calculated as:

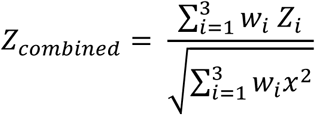

The resulting combined Signed Z-score represented the integrated transcriptional signal of the three related *menF*-associated comparisons. This combined signature was then treated as one independent study-level input in the principal cross-study analysis.

### 2.7. Robust Rank Aggregation meta-analysis

Robust Rank Aggregation (RRA) (Kolde et al., 2012) was used to identify genes showing reproducibly high ranks across the independent study-level inputs. The final meta-analysis consisted of seven ranked gene lists: the combined *menF*-associated signature and the six independent datasets GSE140478, GSE158959, GSE17276, GSE206196, GSE33147, and GSE316857.

Positive and negative transcriptional responses were analyzed independently. For the upregulated analysis, genes with positive signed Z-scores were ranked in descending order, whereas genes with negative signed Z-scores were ranked in ascending order for the downregulated analysis, ensuring that genes with the strongest evidence of upregulation or downregulation were assigned the highest ranks within their respective lists. This procedure generated separate ranked lists representing putatively conserved upregulation and downregulation across the evolutionary experiments. A common gene universe was defined from the union of genes present across the positive and negative ranked lists. Rank matrices were then constructed using the rankMatrix function, and RRA was performed using the aggregateRanks function with the RRA method.

The RRA score represents the probability of observing the observed rank positions of a gene across the input lists under a random-ranking model, with lower scores indicating stronger evidence of consistent high ranking across studies (Kolde et al., 2012). Genes with an RRA score below 0.01 were initially retained as candidate conserved upregulated or downregulated genes. Benjamini–Hochberg-adjusted values were additionally calculated for reporting and assessment of multiple testing (Benjamini & Hochberg, 1995), although the prespecified primary selection criterion for the conserved signature was the RRA score threshold of 0.01.

### 2.8. Directional consistency and recurrence filtering

To ensure that the final conserved signature represented directionally consistent transcriptional adaptation, genes identified simultaneously in the upregulated and downregulated RRA sets were classified as conflicting genes and removed from both sets (after primary selection). The remaining genes were combined into a single core-gene table with an assigned direction of regulation.

The recurrence of each candidate gene was then quantified across the seven independent meta-analysis inputs. For this purpose, each gene was counted once per study-level input in which it was present. The combined *menF*-associated signature was treated as one independent input, consistent with its prior integration. Genes supported by at least six of the seven independent inputs were retained as the final conserved core set. This recurrence criterion was intentionally stringent and was used to prioritize genes whose presence and directional ranking were reproducible across most of the independent evolutionary contexts rather than genes driven by only a subset of studies. The resulting upregulated and downregulated core-gene sets were exported for functional and network analyses.

### 2.6. Functional enrichment analysis

The final conserved gene set was subjected to functional enrichment analysis to determine whether recurrently altered genes were concentrated in specific biological processes or pathways. Gene Ontology (GO) Biological Process enrichment and Kyoto Encyclopedia of Genes and Genomes (KEGG) pathway enrichment were performed using the conserved core genes as the input gene set (Carbon et al., 2021; Kanehisa et al., 2023; Wu et al., 2021). Functional enrichment analysis was performed to characterize the biological functions represented among the conserved genes. Gene Ontology and pathway enrichment analyses were used to identify biological processes and functional categories associated with the conserved transcriptional response.

Functional interpretation was performed at the level of biological processes and pathways rather than individual genes alone. Particular attention was given to terms representing coordinated cellular functions, including motility, chemotaxis, flagellar assembly, transport, metabolism, and surface-associated processes.

### 2.10. Protein–protein interaction and hub-gene analysis

To explore the functional relationships among the conserved recurrent genes, a protein–protein interaction (PPI) network was constructed using the STRING database (version 12.0) (Szklarczyk et al., 2023), with *Escherichia coli* K-12 as the reference organism. Conserved recurrent genes identified by Robust Rank Aggregation (RRA) were mapped to *E. coli* K-12 MG1655 locus tags using the corresponding reference GFF annotation and subsequently submitted to STRING to retrieve protein interaction information. The resulting PPI network was exported from STRING and imported into Cytoscape (version 3.10.4) (Shannon et al., 2003) for network visualization and downstream analysis. Hub genes were identified using the cytoHubba plugin (Chin et al., 2014) in Cytoscape; however, the maximal clique centrality (MCC) method produced identical scores for multiple highly connected nodes and therefore did not provide sufficient resolution for reliable ranking. Consequently, the Degree algorithm was used for final hub-gene prioritization, with genes ranked according to the number of their direct protein–protein interaction partners. Genes were ranked by their degree scores, and the top 20 highest-degree genes were retained as the principal hub-gene set for subsequent analysis. The resulting hub-gene network was visualized with node size proportional to degree, while node colors were used to indicate manually curated functional categories, including flagellar assembly, flagellar motor, chemotaxis, biofilm/curli, regulatory functions, and other functions.

### 2.11. MCODE module detection

To resolve higher-order organization within the interaction network, densely connected sub-networks were identified using the Molecular Complex Detection (MCODE) algorithm within the Cytoscape platform (Bader & Hogue, 2003). MCODE clusters were evaluated according to their network connectivity and subsequently interpreted using their constituent genes and functional annotations. For the principal network visualization, modules were selected according to their biological relevance to the conserved transcriptional signature rather than MCODE score alone. The main figure therefore focused on the functionally interpretable modules representing flagellar/chemotaxis, transport, and curli/biofilm-associated functions. The complete MCODE output was retained for supplementary reporting.

### 2.12. Cross-study hub-gene expression analysis

To evaluate the reproducibility of hub-gene regulation across individual evolutionary experiments, log2 fold-change values for the identified hub genes were extracted from the corresponding study-specific analysis files. The datasets included the two GSE224889 comparisons, GSE122779, GSE140478, GSE158959, GSE17276, GSE206196, GSE33147, and GSE316857. For GSE17276, the input values corresponded to pre-calculated log2 fold changes representing evolved populations relative to their corresponding ancestor [log2(Evolved/Ancestor)] and were used as reported in the original dataset. Gene-level log2 fold changes were subsequently merged across datasets. To facilitate comparison of regulatory patterns across independent evolutionary contexts, the resulting matrix was standardized independently for each gene across comparisons using row-wise Z-score normalization. Thus, the resulting values and color scale represent relative expression patterns within each gene across comparisons rather than directly comparable absolute effect magnitudes across datasets. Hierarchical clustering was then applied to both genes and comparisons, and the resulting patterns were visualized as a heatmap. Functional annotations were displayed alongside the genes to facilitate interpretation of recurrent regulatory patterns across evolutionary experiments.

### 2.13. Data visualization and reproducibility

All data preprocessing, statistical analyses, meta-analysis procedures, and visualizations were performed using R statistical software (version 4.3.3), Cytoscape (version 3.10.4), and the STRING database (version 12.0), as appropriate to the specific analysis. Differential-expression analyses were performed using DESeq2 for datasets with raw RNA-seq count matrices and limma for processed continuous-expression data. Where raw microarray CEL files were available, preprocessing and Robust Multi-array Average (RMA) normalization were performed using the affy package. Cross-study meta-analysis was conducted using the RobustRankAggreg package (Kolde et al., 2012), while protein–protein interaction network construction and hub-gene analysis incorporated STRING-derived interaction information and Cytoscape-based network analysis. UpSet visualization was used (Conway et al., 2017) to display the distribution and recurrence of genes across the seven study-level meta-analysis inputs. Conserved genes identified by Robust Rank Aggregation (RRA) were subsequently used to construct a protein–protein interaction network, from which hub genes were prioritized using the Degree algorithm implemented in cytoHubba in Cytoscape. The top 20 genes ranked by Degree were retained for hub-gene network visualization and downstream functional interpretation. All intermediate study-specific differential-expression tables, Signed Z-score matrices, RRA outputs, conserved-gene tables, locus-tag mappings, and downstream network-analysis files were retained as separate analysis outputs. The complete computational workflow was organized into dataset-specific preprocessing and differential-expression scripts, followed by dedicated scripts for meta-analysis, STRING/network analysis, and figure generation, thereby allowing each stage of the analysis to be independently reproduced and audited.

### 2.14. Use of Artificial Intelligence Tools

OpenAI’s ChatGPT was used as a research assistance tool during the preparation of this study, including support with bioinformatics workflow development, computational coding, data analysis and interpretation, manuscript organization, and language editing. AI-assisted outputs were reviewed, evaluated, and verified by the author. The author retains full responsibility for the accuracy, integrity, reproducibility, and scientific conclusions of the study.

## 3. Results

### 3.1. Cross-study integration reveals recurrent transcriptional responses across diverse *E. coli* K-12 evolutionary experiments

To determine whether distinct adaptive laboratory evolution (ALE) experiments converge on recurrent transcriptional responses, we integrated seven study-level transcriptomic inputs representing diverse evolutionary conditions in *Escherichia coli* K-12. The final inputs comprised a combined *menF*-associated signature derived from three related comparisons and six additional datasets representing thermal adaptation, isoprenol adaptation, long-term glucose limitation, homoserine-associated adaptation, glycerol-associated evolution, and paraquat-associated evolution. The three related *menF*-associated comparisons were consolidated into a single study-level signature before cross-study integration, preventing related comparisons from disproportionately contributing to the meta-analysis while retaining their shared transcriptional signal.

RRA was applied independently to the upregulated and downregulated gene rankings to identify transcriptional responses that recurred across the independent evolutionary contexts. Using an RRA score threshold of <0.01 followed by removal of genes with conflicting directional assignments, we identified genes showing consistent evidence of regulation across the study-level inputs. Applying the predefined recurrence criterion of representation in at least six of the seven inputs yielded 109 conserved core genes, comprising 32 consistently upregulated and 77 consistently downregulated genes. Thus, the conserved signature was predominantly characterized by recurrent transcriptional downregulation, with downregulated genes accounting for approximately 71% of the final core set. The broad recurrence of these genes across otherwise distinct evolutionary experiments indicates that the identified transcriptional signature was not restricted to a single evolutionary trajectory or selective condition, but instead captured transcriptional responses repeatedly associated with adaptation across the examined ALE contexts. The distribution and overlap of genes across the seven study-level inputs are shown in the UpSet plot (Figure 1), highlighting the extent of transcriptional recurrence underlying the final conserved signature.

**Figure 1.**
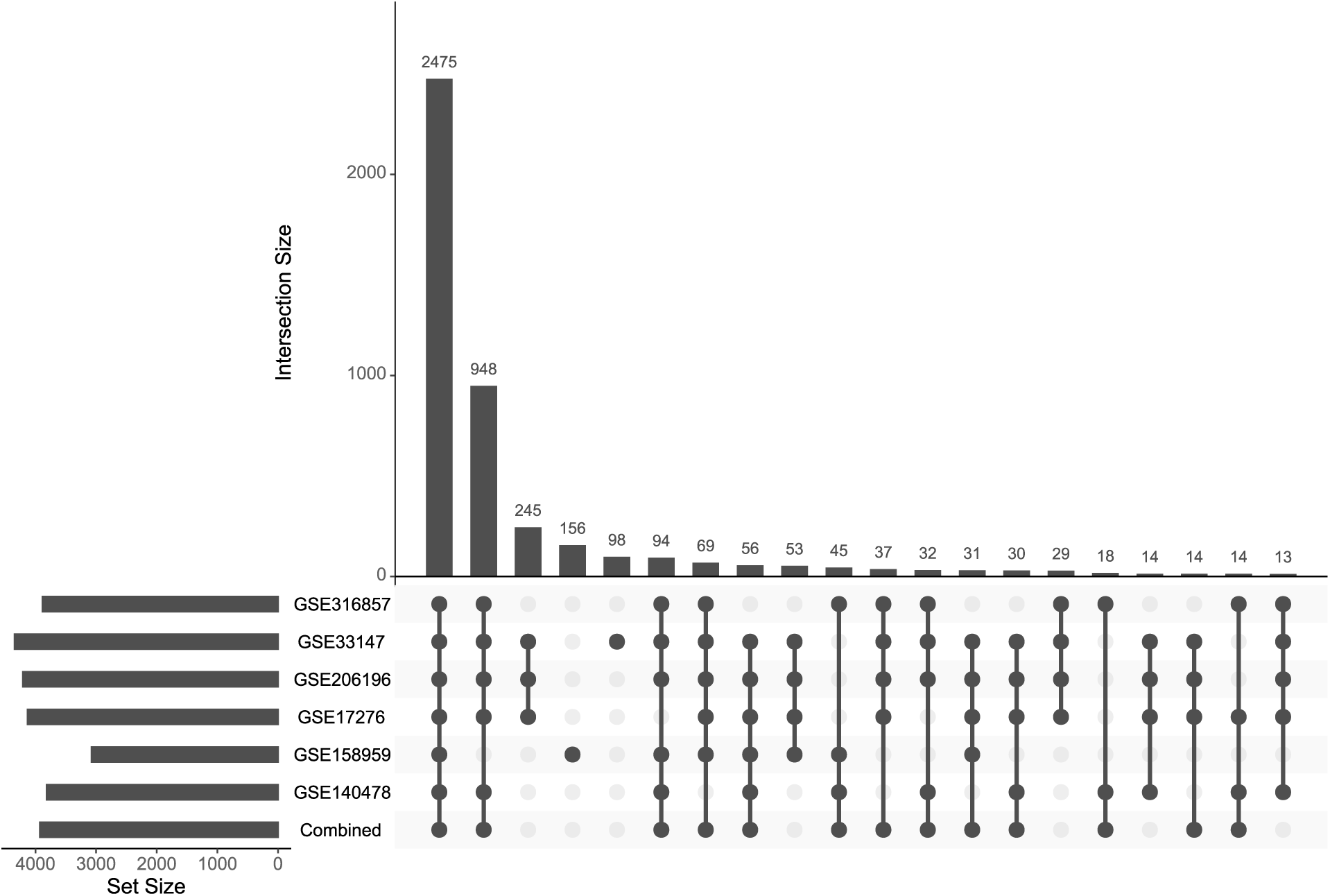
Cross-study recurrence of transcriptional responses across independent *E. coli* K-12 ALE experiments. UpSet plot showing the occurrence and overlap of genes across the seven study-level inputs used in the RRA meta-analysis: the combined *menF*-associated signature and the six independent datasets GSE140478, GSE158959, GSE17276, GSE206196, GSE33147, and GSE316857. Intersections represent shared gene occurrence across study-level inputs, illustrating the extent of transcriptional overlap used to define recurrent adaptive responses.

### 3.2. RRA identifies distinct recurrent upregulated and downregulated gene sets

The directional structure of the conserved transcriptional signature was further examined by ranking genes according to their RRA scores. The top 20 genes in each direction are presented in Figure 2. Among the recurrently upregulated genes, *ftnB* and *pck* showed the strongest RRA scores, followed by *cysU*, *tsuA*, *gcvT*, *gntT*, *yqhD*, and *cfa*. The remainder of the top-ranked upregulated set included *glnK*, *nac*, *cysP*, *nadB*, *aspA*, *ptsG*, *lysA*, *alaE*, *mglA*, *pabC*, *rpsT*, and *rbsC*. Notably, 18 of these 20 genes were recurrent in all seven study-level inputs, while *cysU*, *tsuA*, and *glnK* were represented in six inputs. Thus, the highest-ranked upregulated responses were supported across most or all evolutionary contexts rather than being driven by a limited subset of experiments.

**Figure 2.**
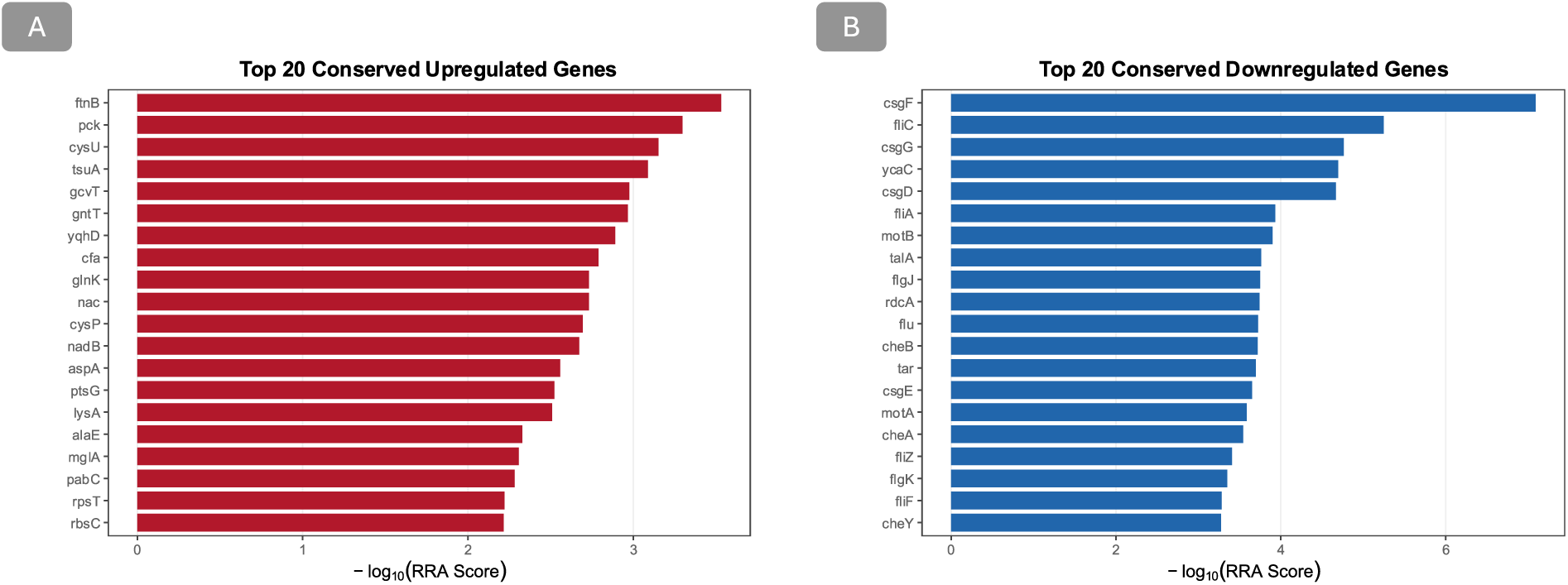
Top recurrent transcriptional responses identified by RRA. (A) The 20 highest-ranked conserved upregulated genes based on the RRA score. (B) The 20 highest-ranked conserved downregulated genes. Bars represent −log10(RRA score), with larger values indicating stronger recurrence across the independent study-level inputs.

Functionally, the recurrently upregulated genes encompassed diverse aspects of cellular physiology, including iron-associated processes (*ftnB*), central carbon metabolism (*pck*), sulfur acquisition (*cysU*, *cysP*), nitrogen regulation (*glnK*, *nac*), carbon and nutrient uptake (*gntT*, *ptsG*, *mglA*), amino-acid metabolism (*aspA*, *lysA*), membrane lipid metabolism (*cfa*), and other metabolic and cellular functions. This functional breadth indicates that recurrent transcriptional activation was not confined to a single metabolic or stress-response pathway, but instead reflected broader physiological remodeling across the independent evolutionary contexts.

In contrast, the downregulated signature showed a pronounced concentration of genes associated with motility, chemotaxis, and surface-associated structures. *csgF* exhibited the lowest RRA score, followed by *fliC*, *csgG*, *ycaC*, *csgD*, *fliA*, *motB*, *talA*, *flgJ*, *rdcA*, *flu*, *cheB*, *tar*, *csgE*, *motA*, *cheA*, *fliZ*, *flgK*, *fliF*, and *cheY*. Eighteen of these 20 genes were represented in all seven study-level inputs, whereas *motB* and *flgJ* were recurrent in six inputs. The recurrence of multiple components from the same cellular systems was particularly notable: flagellar structural and regulatory genes, including *fliC*, *fliA*, *flgJ*, *flgK*, *fliF*, and *fliZ*, occurred together with chemotaxis-associated genes such as *cheA*, *cheB*, *cheY*, and *tar*, while *csgD*, *csgE*, and *csgF* represented the curli-associated program. The complete ranked gene lists, generated figures, and intermediate analysis outputs generated during this study are publicly available in the associated GitHub repository: https://github.com/MJ-Golmohammadi/ecoli-ale-transcriptomic-meta-analysis

### 3.3. Functional enrichment highlights motility and chemotaxis as dominant conserved adaptive programs

To determine whether the conserved genes represented coherent biological processes, the 109-gene core signature was subjected to functional enrichment analysis (Figure 3). The strongest enrichment was observed for biological processes related to cellular movement and flagellum-dependent motility. The most prominent GO Biological Process term was cell motility, which included 23 of the 68 genes annotated to this process and showed an enrichment significance of approximately 3.86 × 10⁻¹⁴. This was followed by bacterial-type flagellum-dependent cell motility, represented by 21 genes, with an FDR of approximately 4.40 × 10⁻¹³. Chemotaxis was also strongly enriched, with 16 conserved genes represented among the annotated background and an FDR of approximately 6.80 × 10⁻¹¹. Additional enriched GO terms reinforced this functional pattern. These included bacterial-type flagellum-dependent swarming motility, regulation of chemotaxis, regulation of locomotion, response to external stimuli, cell projection organization, and cell projection assembly. Several flagellum-related structural terms were also represented, including bacterial-type flagellum and bacterial-type flagellum basal body. Collectively, these enrichment results indicate that the recurrence observed at the gene level was organized at the level of an interconnected physiological system involving environmental sensing, chemotactic signaling, flagellar assembly, and cellular movement rather than reflecting unrelated individual genes.

**Figure 3.**
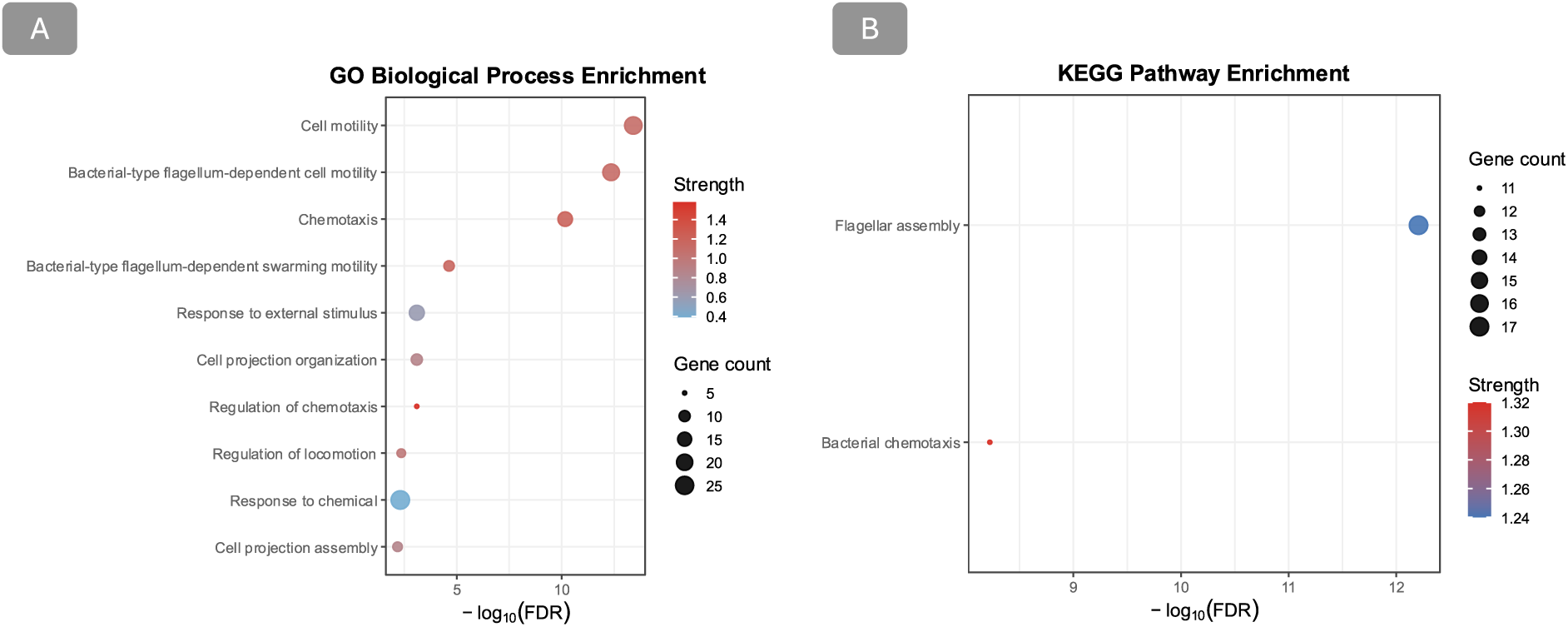
Functional enrichment of the conserved 10G-gene adaptive signature. (A) GO Biological Process enrichment showing the ten most significant enriched terms ranked by false discovery rate. Point size represents the observed gene count, while the x-axis represents −log10(FDR). (B) KEGG pathway enrichment showing the identified pathways associated with the conserved genes, including flagellar assembly and bacterial chemotaxis.

The pathway-level analysis provided an independent representation of the same biological signal. Two KEGG pathways were detected: flagellar assembly, containing 17 of the 37 genes annotated to the pathway, and bacterial chemotaxis, containing 11 of 20 annotated genes. The agreement between the GO and KEGG analyses is particularly informative because both approaches independently identify motility and chemotaxis as the dominant functional components of the conserved transcriptional signature.

The enrichment pattern therefore provides functional context for the strong representation of *fli*, *flg*, *mot*, and *che* genes in the RRA results. Rather than representing independent responses of individual genes, their recurrence suggests coordinated remodeling of a cellular system that couples environmental sensing to locomotion. The simultaneous recurrence of multiple flagellar structural components, motor proteins, and chemotaxis regulators is especially consistent with a system-level transcriptional shift in which the investment in motility-associated functions is repeatedly reduced during adaptation.

### 3.4. Network analysis reveals a highly connected adaptive core dominated by motility and chemotaxis

Because functional enrichment indicated strong representation of motility and chemotaxis genes, the conserved gene set was further examined in the context of protein–protein interactions. The STRING network constructed from the complete 109-gene core signature contained 109 nodes and 581 edges, with an average node degree of 10.7 and an average local clustering coefficient of 0.656. The observed network contained substantially more interactions than expected by chance, with an expected number of approximately 137 edges and a PPI enrichment *P* value of <1.0 × 10⁻¹⁶.

The network structure was characterized by a prominent interconnected motility/chemotaxis component (Figure 4). Degree-based hub prioritization identified *fliC* as the highest-degree node, with a degree score of 33. *evgS* and *fliA* followed with degree scores of 30, while *cheA*, *cheB*, and *cheW* each had a degree of 29. Additional high-degree nodes included *fliD*, *motA*, *motB*, *flgG*, *flgK*, *cheY*, *tar*, and *fliF*.

**Figure 4.**
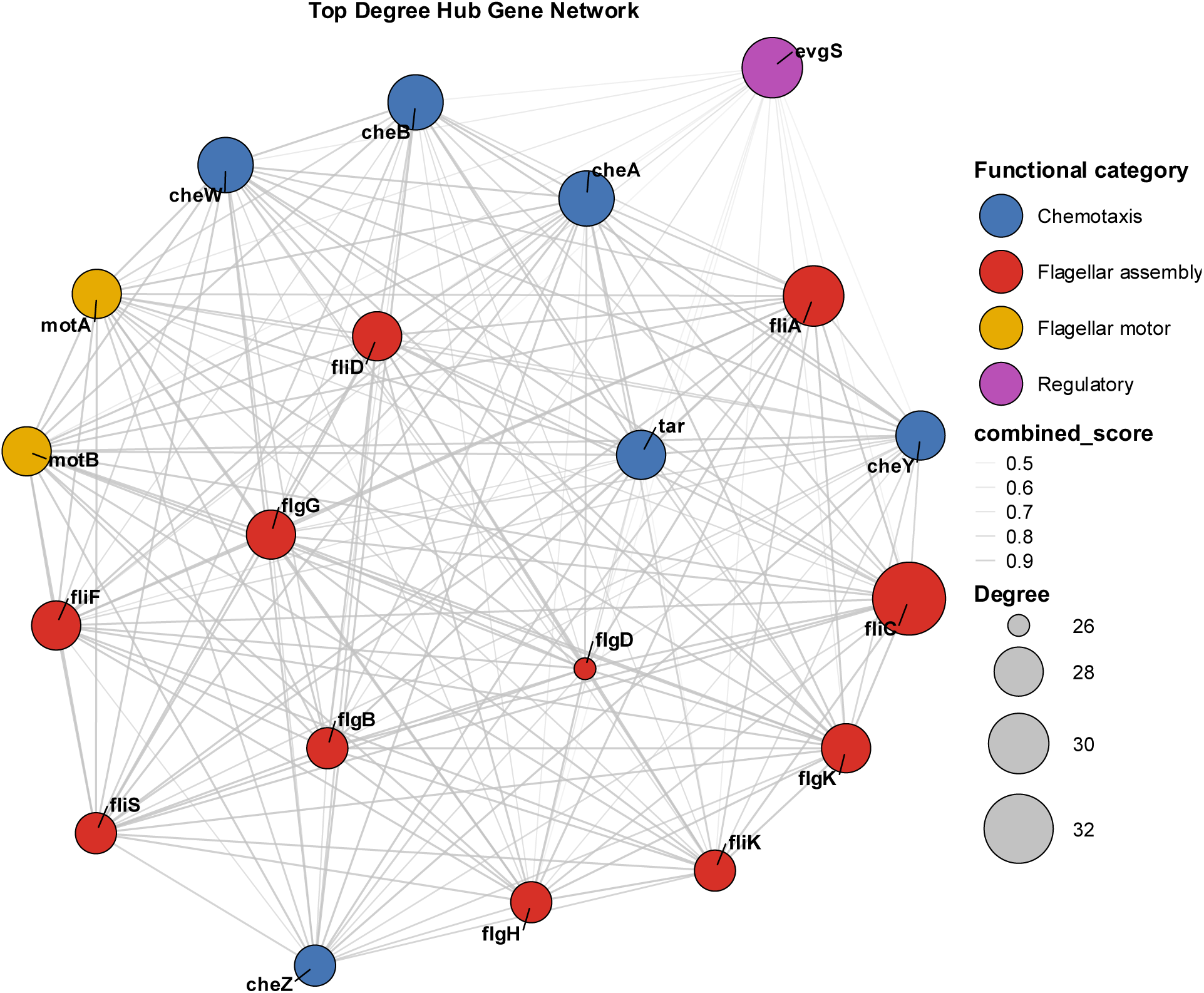
Degree-based hub network of the conserved adaptive signature. Protein–protein interaction network constructed from the top 20 genes ranked by degree. Node size represents degree, while node categories distinguish flagellar assembly, flagellar motor, chemotaxis, and regulatory.

This network organization substantially strengthens the interpretation obtained from RRA and enrichment analyses. *fliC*, which encodes the major flagellin structural component, was simultaneously the highest-degree network hub and one of the most strongly recurrent downregulated genes in the meta-analysis. Likewise, *fliA*, *motA*, *motB*, *cheA*, *cheB*, and *cheY* were represented among the most connected nodes while also contributing to the recurrent downregulated signal. The convergence of independent analytical layers—rank recurrence, functional enrichment, and network connectivity—therefore identifies the flagellar/chemotaxis system as a central component of the conserved adaptive response. The hub network also contained a distinct group of curli/biofilm-associated genes. *csgA*, *csgD*, *csgE*, *csgF*, *csgG*, and *flu* were classified within the biofilm/curli category, while *evgS* represented a regulatory category in the network visualization. The presence of these genes alongside the dominant motility and chemotaxis hubs indicates that the conserved adaptive signature extends beyond locomotion to include remodeling of surface-associated and regulatory functions.

### 3.5. Network modules reveal coordinated flagellar, transport, and curli-associated structures

To identify discrete functional modules within the conserved PPI network, MCODE clustering identified six modules. Cluster 1 was the largest and most densely connected module (27 nodes, 345 edges, score = 26.538) and was dominated by flagellar and chemotaxis genes, including structural, motor, and signaling components. This extensive connectivity was consistent with the enrichment and hub-gene analyses, further supporting coordinated remodeling of motility and environmental sensing. Cluster 2 contained ten genes (44 edges, score = 9.778) but showed a relatively heterogeneous functional composition spanning stress-associated and metabolic functions; therefore, despite its high MCODE score, it was not included in the final visualization. In contrast, Cluster 3 comprised eight genes (*cysP*, *malE*, *modF*, *cysU*, *malF*, *btuD*, *mglA*, and *rbsC*; 28 edges, score = 8.0) and was primarily associated with transport and nutrient acquisition. Cluster 4 contained five curli-associated genes (*csgD*, *csgA*, *csgF*, *csgE*, and *csgG*; 10 edges, score = 5.0), representing a compact but highly coherent surface-associated module. The biologically interpretable modules selected for visualization are shown in Figure 5. Collectively, these modules comprised a dominant flagellar/chemotaxis module, a transport and nutrient-acquisition module, and a curli-associated module, indicating recurrent remodeling of motility, resource acquisition, and surface-associated functions across the evolutionary contexts.

**Figure 5.**
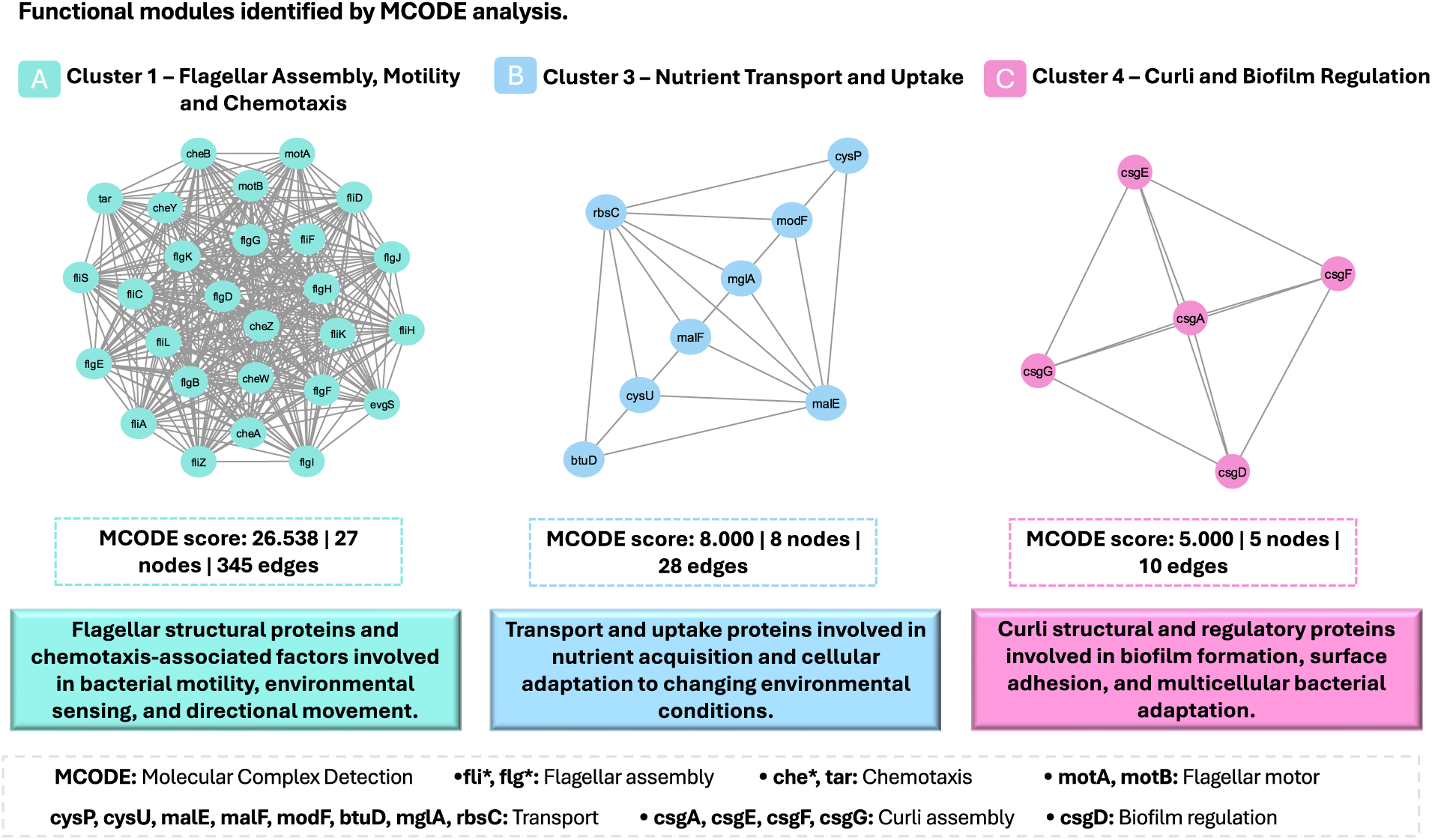
Functionally coherent modules within the conserved adaptive interaction network. MCODE analysis of the STRING network identified multiple densely connected modules. Three biologically coherent modules are shown: Cluster 1, representing flagellar and chemotaxis functions; Cluster 3, representing transport and nutrient acquisition; and Cluster 4, representing the curli-associated system. Cluster 2 was excluded from the visualization because, despite its higher MCODE score, its genes showed comparatively heterogeneous functional composition.

### 3.6. Cross-study expression profiles support recurrent remodeling of hub-associated systems

To examine the reproducibility of hub-gene regulation across individual evolutionary contexts, study-specific log2 fold changes were compiled and visualized by hierarchical clustering (Figure 6). The resulting heatmap revealed a consistent pattern of relative downregulation among the flagellar and chemotaxis-associated hubs, including *fliC*, *fliA*, *motA*, *motB*, *cheA*, *cheB*, *cheY*, and multiple *fli* and *flg* genes, together with the curli-associated genes *csgA*, *csgD*, *csgE*, *csgF*, and *csgG*. These patterns were consistent with the directionally filtered RRA results and the MCODE modules, further supporting the recurrence of motility-, chemotaxis-, and curli-associated transcriptional attenuation across evolutionary contexts. Importantly, the heatmap captures relative expression patterns rather than directly comparable absolute effect magnitudes across heterogeneous datasets; for GSE17276, the corresponding values represented reported log2(Evolved/Ancestor) expression ratios. Overall, the hub-gene profiles indicate that conserved adaptation was characterized by recurrent changes in the same functional systems across experiments, although the magnitude of individual gene-level responses varied between evolutionary contexts.

**Figure 6.**
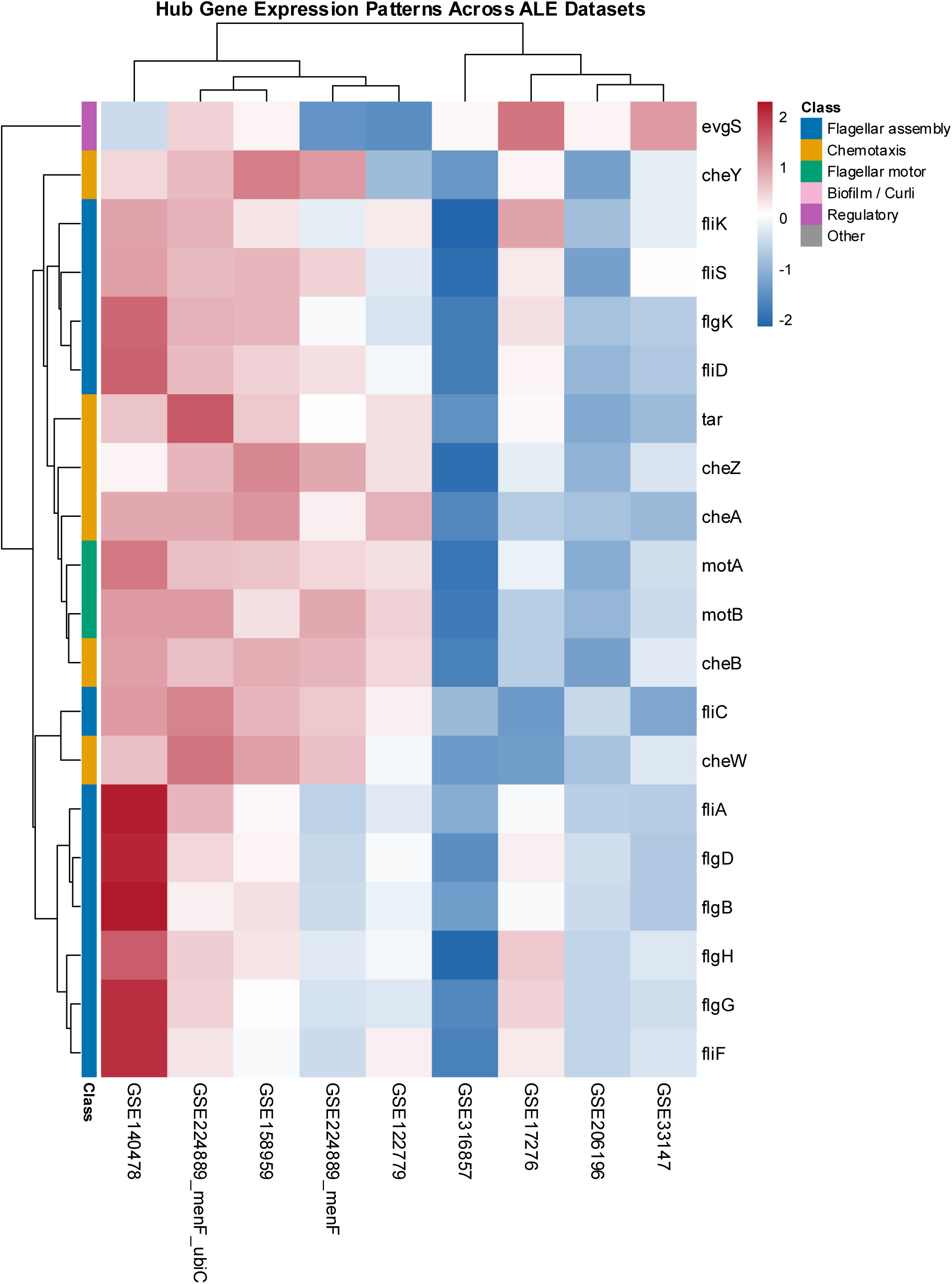
Cross-study expression patterns of conserved hub genes. Hierarchical clustering heatmap showing standardized differential-expression effect estimates for the hub genes across the individual ALE comparisons. Values were standardized independently for each gene across comparisons; therefore, color represents relative expression patterns among evolutionary contexts rather than absolute effect magnitudes. For GSE17276, the input values correspond to pre-calculated log2(Evolved/Ancestor) expression ratios. Functional annotations identify flagellar assembly, chemotaxis, flagellar motor, biofilm/curli, regulatory, and other hub-gene categories.

## 4. Discussion

Adaptive laboratory evolution provides a powerful experimental framework for investigating how microbial populations respond to sustained genetic and environmental perturbations. Although parallel evolution can produce reproducible phenotypic outcomes, the molecular routes leading to those outcomes are often highly variable (Barrick et al., 2009; Fong et al., 2005; Travisano et al., 1995). Different evolved populations may acquire distinct mutations, alter different regulatory components, or redistribute cellular resources through partially different physiological pathways (Elena & Lenski, 2003). This variability creates a central challenge for transcriptomic studies of experimental evolution: responses detected in an individual experiment may reflect the specific selective environment, genetic background, or evolutionary trajectory rather than a general feature of adaptation (X. Wang et al., 2018). Previous multi-omics analysis of experimentally evolved *E. coli* has shown that transcriptomic adaptations can be strongly conserved across genetically diverse strains, despite substantial differences in their genomes and underlying causal mutations (Kavvas et al., 2022). These findings indicate that evolutionary convergence can occur at the level of systems-wide physiological and transcriptional states rather than requiring identical genetic changes. Such evidence supports comparative approaches that integrate transcriptomic responses across independent evolutionary experiments to identify conserved features of adaptation. By integrating seven study-level transcriptomic inputs representing diverse evolutionary contexts in *Escherichia coli* K-12, the present study addressed this problem by searching for transcriptional responses that recur across independent ALE experiments. The identification of 109 conserved genes, including 32 upregulated and 77 downregulated genes, provides evidence that despite substantial context dependence, a measurable component of transcriptional remodeling is repeatedly associated with laboratory adaptation.

### 4.1. Conserved transcriptional adaptation is dominated by recurrent attenuation rather than uniform induction

One of the most notable features of the conserved signature was its directional asymmetry. The final core consisted of substantially more downregulated genes than upregulated genes, with 77 genes showing recurrent negative responses compared with 32 recurrent positive responses. This pattern suggests that a major component of the conserved transcriptional response across the examined ALE experiments is not the universal activation of stress-protective pathways, but rather the repeated attenuation of particular cellular functions. Similar observations from previous experimental evolution studies indicate that adaptation can involve extensive remodeling of transcriptional programs and redistribution of cellular functions rather than a uniform induction of stress-associated responses (Fong et al., 2005; Kavvas et al., 2022).

Reducing the expression of functions that are less beneficial under a particular environment may allow cells to allocate resources toward processes that are more important for growth and survival (Basan et al., 2015). In this context, recurrent transcriptional attenuation does not necessarily indicate loss of cellular function (Kavvas et al., 2022); instead, it may represent a common adaptive strategy across different evolutionary conditions. The strong recurrence of motility-, chemotaxis-, and surface-associated genes within the downregulated signature supports this interpretation, as these functions were repeatedly reduced across otherwise distinct evolutionary experiments.

In contrast, the recurrently upregulated genes represented a broader and more heterogeneous set of metabolic, transport, biosynthetic, and regulatory functions, including *cysU*, *cysP*, *gcvT*, *gntT*, *glnK*, *nac*, *ptsG*, *lysA*, *mglA*, and *ompF*. These genes did not form a single dominant pathway, but instead represented diverse functional processes, suggesting that recurrent upregulation involved broad remodeling of cellular physiology rather than activation of a single conserved pathway. Taken together, the different patterns of up- and downregulation argue against a stereotyped universal stress response in which the same stress-protective genes are activated under all conditions. Instead, the results support a model of conserved adaptation in which some cellular programs are repeatedly attenuated, while metabolic and regulatory functions are adjusted more flexibly according to environmental demands (Dalldorf, Rychel, et al., 2024; Ferenci, 2005; Fong et al., 2006; Kavvas et al., 2022).

### 4.2. Flagellar and chemotaxis functions emerge as the central conserved adaptive system

The most consistent feature of the conserved transcriptional signature was the recurrent attenuation of flagellar and chemotaxis functions. This pattern was supported across complementary analytical levels, with multiple flagellar and chemotaxis genes—including *fliC*, *fliA*, *flgJ*, *motA*, *motB*, *cheA*, *cheB*, *cheW*, *cheY*, and *tar*—among the recurrently downregulated genes, while functional enrichment and KEGG analysis independently highlighted motility, bacterial chemotaxis, and flagellar assembly. Network analysis further showed that these genes were embedded within a densely connected interaction structure, with the largest MCODE module comprising 27 nodes and 345 edges and being dominated by flagellar and chemotaxis components. Together, these findings indicate that the conserved signal involved coordinated attenuation of an interconnected motility system rather than isolated changes in individual genes. One possible explanation is a trade-off between the fitness benefits of motility and the cellular cost of maintaining the flagellar apparatus. Although motility and chemotaxis can be beneficial for environmental exploration, flagellar biosynthesis and activity require substantial cellular resources. In well-mixed laboratory environments, where dispersal and active nutrient searching may provide limited additional benefits, selection may therefore favor reduced investment in motility-associated functions during prolonged adaptation (Lisevich et al., 2025; Schwartz et al., 2025). In addition, evolutionary trajectory of motility depends strongly on the selective environment: selection for migration can favor increased swimming capacity, whereas the balance between motility and growth can shift substantially with environmental conditions (Fraebel et al., 2017).

### 4.3. Recurrent chemotaxis suppression is associated with coordinated remodeling of environmental sensing and motility

The recurrent downregulation of chemotaxis genes alongside flagellar components suggests that the conserved response extended beyond the structural machinery of movement to the sensory and signaling system that regulates motility. Chemotaxis enables *E. coli* to detect and exploit chemical gradients, but its fitness benefit is strongly dependent on environmental conditions, nutrient availability, and the potential to encounter exploitable gradients. Experimental studies have shown that *E. coli* adjusts its investment in motility according to the expected benefit of chemotaxis, with both the cost of maintaining flagellar machinery and the fitness benefit of chemotactic behavior varying with nutritional conditions (Lisevich et al., 2025; Ni et al., 2020). Importantly, chemotaxis can remain beneficial even in initially homogeneous cultures because bacterial growth, nutrient consumption, and metabolite secretion can generate local chemical gradients that subsequently influence cell redistribution and nutrient acquisition (Ni et al., 2020). Thus, the recurrent attenuation observed in the present study may reflect a broader adaptive pattern (Lisevich et al., 2025) that is not restricted to the nutritionally homogeneous conditions typically used in laboratory evolution. The recurrence of *cheA*, *cheB*, *cheW*, *cheY*, and *tar* across the nine comparisons, which encompassed distinct evolutionary environments, suggests that remodeling of chemotactic sensing and signaling accompanied adaptation across a broad range of selective conditions. The coordinated recurrence of chemotaxis and flagellar genes in our meta-analysis is notable because it was observed despite substantial differences among the evolutionary comparisons. The organization of these genes within the same densely connected interaction module further supports the interpretation that the recurrent signal represents coordinated remodeling of an integrated motility–chemotaxis system rather than independent changes in individual genes. Nevertheless, because our meta-analysis identifies recurrent transcriptional patterns rather than directly measuring fitness effects, the observed attenuation does not by itself demonstrate that reduced chemotaxis was beneficial under all evolutionary conditions. Instead, it identifies remodeling of the motility–chemotaxis system as a recurrent feature of transcriptional adaptation across the diverse evolutionary contexts represented in this study, providing a basis for future experimental work to determine when and how changes in this system contribute causally to fitness.

### 4.4 Recurrent attenuation of curli-associated functions suggests broader surface-interaction remodeling

The recurrent downregulation of the curli-associated genes *csgD*, *csgA*, *csgE*, *csgF*, and *csgG* identifies a second functionally coherent component of the conserved adaptive response. These genes form the core of the curli regulatory and assembly program, with *csgD* serving as a major regulator of *csgBA* expression and *csgE/F/G* contributing to curli assembly and export (Barnhart & Chapman, 2006; Prigent-Combaret et al., 2001). The coordinated recurrence of these genes in the downregulated signature, together with their complete connectivity within the five-gene network module, therefore suggests remodeling of the curli-associated program rather than independent changes in individual structural genes. Curli are extracellular amyloid structures that contribute to surface adhesion and biofilm matrix formation, but their expression is highly dependent on environmental and physiological conditions, including temperature, osmolarity, nutrient availability, oxygen status, and growth state (Barnhart & Chapman, 2006; Gualdi et al., 2007; Hufnagel et al., 2015).

The simultaneous attenuation of curli-associated and flagellar/chemotaxis genes is particularly noteworthy. In *E. coli*, these systems are functionally interconnected through regulatory networks that coordinate motility with the transition toward surface-associated growth. CsgD promotes curli and extracellular matrix production while repressing components of the flagellar system, and c-di-GMP provides an additional regulatory link between motility and surface-associated functions (Guttenplan & Kearns, 2013; Ogasawara et al., 2011; Thomason et al., 2012). However, the pattern observed in our meta-analysis should not be interpreted simply as a switch from a motile to a sessile state (Ogasawara et al., 2011), because both motility/chemotaxis and curli-associated genes were recurrently downregulated across the evolutionary comparisons. Instead, their concurrent attenuation suggests a broader remodeling of cellular functions involved in environmental interaction and surface-associated behavior (Rossi et al., 2018). Both motility and curli production require coordinated expression of multiple genes and substantial cellular investment, while their fitness value is strongly dependent on environmental context (Barnhart & Chapman, 2006; Evans & Chapman, 2014; Pesavento et al., 2008; Rossi et al., 2018).

This interpretation also provides a more cautious explanation for the apparent divergence from the classical motile–sessile regulatory relationship. Rather than indicating a uniform transition toward either motility or surface attachment (Serra & Hengge, 2021), the recurrent reduction of both programs may reflect decreased investment in multiple extracellular functions during adaptation, accompanied by greater emphasis on other physiological processes such as nutrient acquisition, transport, metabolism, and cellular maintenance. Importantly, the recurrence of *csgD* and its associated genes does not by itself demonstrate reduced biofilm formation (Serra & Hengge, 2021; Tursi & Tükel, 2018), because curli expression is highly condition-dependent and *E. coli* can employ alternative mechanisms of surface adhesion and biofilm formation (Klemm & Schembri, 2004; X. Wang et al., 2004). Thus, the most conservative interpretation of our findings is that curli-associated functions represent a second recurrently remodeled cellular system during ALE, complementing the dominant attenuation of flagellar and chemotaxis functions. The convergence of these two modules suggests that adaptation across the diverse evolutionary environments examined here involved coordinated remodeling of cellular investment in environmental interaction, rather than representing a simple switch between motile and sessile states.

### 4.5. A recurrent transport signature indicates remodeling of resource acquisition

The transport-associated module provides a complementary dimension of the conserved adaptive response, comprising *cysP*, *malE*, *modF*, *cysU*, *malF*, *btuD*, *mglA*, and *rbsC*, which are involved in the uptake of diverse nutrients and metabolites. Although smaller and less densely connected than the dominant flagellar/chemotaxis module, this module indicates that recurrent adaptation also involved resource acquisition. Consistent with this network-level pattern, several transport- and metabolism-associated genes, including *cysU*, *tsuA*, *gntT*, *ptsG*, *mglA*, *rbsC*, *ompF*, and *lysP*, were recurrently upregulated by RRA (Carreón-Rodríguez et al., 2023; Zhong et al., 2009). Together, these findings suggest recurrent remodeling of nutrient acquisition across the evolutionary environments examined in the present study.

Transport and metabolic remodeling can directly influence bacterial fitness because nutrient uptake determines access to cellular resources, while the regulation of uptake systems is closely coupled to metabolic demand and environmental nutrient availability (Okano et al., 2019). In *E. coli*, transporter expression is extensively regulated according to nutrient conditions and the physiological state of the cell, allowing uptake capacity to be adjusted to changing metabolic requirements (van Heeswijk et al., 2013). The coexistence of recurrently altered transport and metabolic genes with strong attenuation of motility-associated programs therefore provides a broader view of resource reorganization during adaptation. Thus, the conserved signature suggests that adaptation repeatedly modifies both what the cell acquires from its environment and which cellular activities it invests in after acquisition (Iyer et al., 2021). Rather than indicating a single universal metabolic strategy, these patterns are more consistent with recurrent reconfiguration of resource acquisition and cellular resource allocation across the distinct evolutionary environments represented in the present study.

### 4.6. Study Limitations, Strengths, and Future Directions

Several considerations should be taken into account when interpreting the present findings. The analyzed datasets were generated using different experimental designs, evolutionary durations, environmental conditions, and transcriptomic platforms, introducing biological and technical heterogeneity that may limit direct comparison of absolute expression changes. Although the rank-based integration strategy reduces the influence of differences in effect-size scales and platform characteristics, it cannot completely eliminate study-specific variation (Kolde et al., 2012). In addition, some evolutionary environments shared broad physiological or selective features, meaning that the observed recurrence may partly reflect common environmental constraints rather than fully independent evolutionary processes. The present analysis was also restricted to transcriptional responses and therefore cannot establish whether recurrent expression changes were directly adaptive, resulted from common genetic alterations, or represented downstream consequences of other evolved phenotypes. Nevertheless, the identification of recurrent transcriptional signatures across multiple independent evolutionary experiments, together with their convergence in functionally coherent networks and modules, represents a major strength of the study and provides evidence that extends beyond findings from any single ALE experiment. Future studies combining longitudinal transcriptomics with whole-genome sequencing, proteomics, metabolomics, and experimentally measured fitness and phenotypic traits within the same evolved populations would help distinguish causal adaptive changes from correlated responses and determine whether the recurrent systems identified here represent general features of bacterial adaptation or are specific to particular laboratory regimes.

## 5. Conclusion

The present cross-study transcriptomic meta-analysis identifies a conserved component of adaptive transcriptional remodeling across diverse *E. coli* K-12 ALE experiments. By integrating seven study-level transcriptomic inputs and applying directional Signed Z-score harmonization, RRA, stringent recurrence filtering, functional enrichment, and protein–protein interaction analysis, we identified 109 recurrent core genes, comprising 32 upregulated and 77 downregulated genes. The predominance of downregulated genes was accompanied by a striking and reproducible concentration of flagellar, chemotaxis, and surface-associated functions.

Multiple independent analytical layers converged on flagellar assembly and chemotaxis as the dominant conserved adaptive programs. Highly connected genes including *fliC*, *fliA*, *cheA*, *cheB*, *cheW*, *motA*, and *motB* formed a central network structure, while a distinct curli-associated module and a transport-associated module revealed additional dimensions of recurrent adaptive remodeling. These findings suggest that adaptation across the heterogeneous laboratory environments represented in this study repeatedly involves remodeling of cellular investment in motility, environmental sensing, surface-associated functions, and resource acquisition.

Importantly, the results do not imply a single universal transcriptional state or identical evolutionary mechanisms across populations. Rather, they demonstrate that molecularly diverse evolutionary trajectories can converge on recurrent physiological systems. The identified genes and modules therefore represent candidate components of a conserved adaptive architecture in *E. coli* K-12. Future integration with genomic, proteomic, metabolic, and experimental fitness data will be necessary to determine the causal mechanisms underlying these recurrent transcriptional changes and to establish whether the identified systems represent general principles of bacterial adaptation beyond the laboratory evolution contexts examined here.

## Acknowledgements

The author gratefully acknowledges the researchers who generated and publicly shared the transcriptomic datasets analyzed in this study through the NCBI Gene Expression Omnibus (GEO). The availability of these datasets enabled their integration across independent evolutionary experiments. The author also acknowledges the developers and maintainers of the open-source computational tools, statistical packages, and biological databases used throughout the analysis, including R, Bioconductor, STRING, and Cytoscape. OpenAI’s ChatGPT was used as an assistive tool during manuscript preparation, language editing, and refinement of the presentation. All analyses, scientific interpretations, and conclusions were independently evaluated and verified by the author.

## Funding

This research was conducted without external financial support.

## Conflict of Interest

The author declares that there are no conflicts of interest or competing interests associated with this work.

## Data Availability Statement

All transcriptomic datasets analyzed in this study are publicly available through the NCBI Gene Expression Omnibus (GEO) under the accession numbers GSE122779, GSE140478, GSE158959, GSE17276, GSE206196, GSE224889, GSE33147, and GSE316857. The analyses were conducted using publicly available computational tools, statistical packages, and biological databases. The processed data, analysis scripts, intermediate results, The processed data, analysis scripts, intermediate results, and figures generated in this study are available in the GitHub repository: https://github.com/MJ-Golmohammadi/ecoli-ale-transcriptomic-meta-analysis

## Author Contributions

Mohammad Javad Golmohammadi conceived and designed the study, performed data acquisition, preprocessing, bioinformatics analyses, statistical analysis, data interpretation, visualization, and manuscript preparation. The author reviewed and approved the final version of the manuscript.

## Notes

https://github.com/MJ-Golmohammadi/ecoli-ale-transcriptomic-meta-analysis

### Competing Interest Statement

The authors have declared no competing interest.

## References

Abdel-Salam, E. M., Figueroa-Gonzalez, T., & Leister, D. (2023). Transcriptomic meta-analysis and functional validation identify genes linked to adaptation and involved in high-light acclimation in Synechocystis sp. PCC 6803. Frontiers in Photobiology, 1. 10.3389/fphbi.2023.1290382

Alkim, C., Farias, D., Fredonnet, J., Serrano-Bataille, H., Herviou, P., Picot, M., Slama, N., Dejean, S., Morin, N., Enjalbert, B., & François, J. M. (2022). Toxic effect and inability of L-homoserine to be a nitrogen source for growth of Escherichia coli resolved by a combination of in vivo evolution engineering and omics analyses. Frontiers in Microbiology, 13. 10.3389/fmicb.2022.1051425

Anand, A., Olson, C. A., Yang, L., Sastry, A. V., Catoiu, E., Choudhary, K. S., Phaneuf, P. V., Sandberg, T. E., Xu, S., Hefner, Y., Szubin, R., Feist, A. M., & Palsson, B. O. (2019). Pseudogene repair driven by selection pressure applied in experimental evolution. Nature Microbiology, 4(3), 386–389. 10.1038/s41564-018-0340-2

Babel, H., & Krömer, J. O. (2020). Evolutionary engineering of E. coli MG1655 for tolerance against isoprenol. Biotechnology for Biofuels, 13(1), 183. 10.1186/s13068-020-01825-6

Bader, G. D., & Hogue, C. W. (2003). An automated method for finding molecular complexes in large protein interaction networks. BMC Bioinformatics, 4(1), 2. 10.1186/1471-2105-4-2

Barnhart, M. M., & Chapman, M. R. (2006). Curli Biogenesis and Function. Annual Review of Microbiology, 60(1), 131–147. 10.1146/annurev.micro.60.080805.142106

Barrick, J. E., Yu, D. S., Yoon, S. H., Jeong, H., Oh, T. K., Schneider, D., Lenski, R. E., & Kim, J. F. (2009). Genome evolution and adaptation in a long-term experiment with Escherichia coli. Nature, 461(7268), 1243–1247. 10.1038/nature08480

Basan, M., Hui, S., Okano, H., Zhang, Z., Shen, Y., Williamson, J. R., & Hwa, T. (2015). Overflow metabolism in Escherichia coli results from efficient proteome allocation. Nature, 528(7580), 99–104. 10.1038/nature15765

Benjamini, Y., & Hochberg, Y. (1995). Controlling the False Discovery Rate: A Practical and Powerful Approach to Multiple Testing. Journal of the Royal Statistical Society Series B: Statistical Methodology, 57(1), 289–300. 10.1111/j.2517-6161.1995.tb02031.x

Carbon, S., Douglass, E., Good, B. M., Unni, D. R., Harris, N. L., Mungall, C. J., Basu, S., Chisholm, R. L., Dodson, R. J., Hartline, E., Fey, P., Thomas, P. D., Albou, L.-P., Ebert, D., Kesling, M. J., Mi, H., Muruganujan, A., Huang, X., Mushayahama, T.,…Elser, J. (2021). The Gene Ontology resource: enriching a GOld mine. Nucleic Acids Research, 49(D1), D325–D334. 10.1093/nar/gkaa1113

Carreón-Rodríguez, O. E., Gosset, G., Escalante, A., & Bolívar, F. (2023). Glucose Transport in Escherichia coli: From Basics to Transport Engineering. Microorganisms, 11(6), 1588. 10.3390/microorganisms11061588

Chen, C., Grennan, K., Badner, J., Zhang, D., Gershon, E., Jin, L., & Liu, C. (2011). Removing Batch Effects in Analysis of Expression Microarray Data: An Evaluation of Six Batch Adjustment Methods. PLoS ONE, 6(2), e17238. 10.1371/journal.pone.0017238

Chin, C.-H., Chen, S.-H., Wu, H.-H., Ho, C.-W., Ko, M.-T., & Lin, C.-Y. (2014). cytoHubba: identifying hub objects and sub-networks from complex interactome. BMC Systems Biology, 8(S4), S11. 10.1186/1752-0509-8-S4-S11

Choe, D., Lee, J. H., Yoo, M., Hwang, S., Sung, B. H., Cho, S., Palsson, B., Kim, S. C., & Cho, B.-K. (2019). Adaptive laboratory evolution of a genome-reduced Escherichia coli. Nature Communications, 10(1), 935. 10.1038/s41467-019-08888-6

Conway, J. R., Lex, A., & Gehlenborg, N. (2017). UpSetR: an R package for the visualization of intersecting sets and their properties. Bioinformatics, 33(18), 2938–2940. 10.1093/bioinformatics/btx364

Cooper, T. F., Rozen, D. E., & Lenski, R. E. (2003). Parallel changes in gene expression after 20,000 generations of evolution in *Escherichia coli*. Proceedings of the National Academy of Sciences, 100(3), 1072–1077. 10.1073/pnas.0334340100

Cronan, J. E. (2014). *Escherichia coli* as an Experimental Organism. In Encyclopedia of Life Sciences. Wiley. 10.1002/9780470015902.a0002026.pub2

Dalldorf, C., Hefner, Y., Szubin, R., Johnsen, J., Mohamed, E., Li, G., Krishnan, J., Feist, A. M., Palsson, B. O., & Zielinski, D. C. (2024). Diversity of Transcriptional Regulatory Adaptation in *E. coli*. Molecular Biology and Evolution, 41(11). 10.1093/molbev/msae240

Dalldorf, C., Rychel, K., Szubin, R., Hefner, Y., Patel, A., Zielinski, D. C., & Palsson, B. O. (2024). The hallmarks of a tradeoff in transcriptomes that balances stress and growth functions. MSystems, 9(7). 10.1128/msystems.00305-24

Dragosits, M., & Mattanovich, D. (2013). Adaptive laboratory evolution – principles and applications for biotechnology. Microbial Cell Factories, 12(1), 64. 10.1186/1475-2859-12-64

Elena, S. F., & Lenski, R. E. (2003). Evolution experiments with microorganisms: the dynamics and genetic bases of adaptation. Nature Reviews Genetics, 4(6), 457–469. 10.1038/nrg1088

Evans, M. L., & Chapman, M. R. (2014). Curli biogenesis: Order out of disorder. Biochimica et Biophysica Acta (BBA) - Molecular Cell Research, 1843(8), 1551–1558. 10.1016/j.bbamcr.2013.09.010

Favate, J. S., Liang, S., Cope, A. L., Yadavalli, S. S., & Shah, P. (2022). The landscape of transcriptional and translational changes over 22 years of bacterial adaptation. ELife, 11. 10.7554/eLife.81979

Ferenci, T. (2005). Maintaining a healthy SPANC balance through regulatory and mutational adaptation. Molecular Microbiology, 57(1), 1–8. 10.1111/j.1365-2958.2005.04649.x

Fong, S. S., Joyce, A. R., & Palsson, B. Ø. (2005). Parallel adaptive evolution cultures of *Escherichia coli* lead to convergent growth phenotypes with different gene expression states. Genome Research, 15(10), 1365–1372. 10.1101/gr.3832305

Fong, S. S., Nanchen, A., Palsson, B. O., & Sauer, U. (2006). Latent Pathway Activation and Increased Pathway Capacity Enable Escherichia coli Adaptation to Loss of Key Metabolic Enzymes. Journal of Biological Chemistry, 281(12), 8024–8033. 10.1074/jbc.M510016200

Fraebel, D. T., Mickalide, H., Schnitkey, D., Merritt, J., Kuhlman, T. E., & Kuehn, S. (2017). Environment determines evolutionary trajectory in a constrained phenotypic space. ELife, c. 10.7554/eLife.24669

Franchini, A. G., & Egli, T. (2006). Global gene expression in Escherichia coli K-12 during short-term and long-term adaptation to glucose-limited continuous culture conditions. Microbiology, 152(7), 2111–2127. 10.1099/mic.0.28939-0

Gautier, L., Cope, L., Bolstad, B. M., & Irizarry, R. A. (2004). affy—analysis of *Affymetrix GeneChip* data at the probe level. Bioinformatics, 20(3), 307–315. 10.1093/bioinformatics/btg405

Gualdi, L., Tagliabue, L., & Landini, P. (2007). Biofilm formation-gene expression relay system in Escherichia coli: modulation of sigmaS-dependent gene expression by the CsgD regulatory protein via sigmaS protein stabilization. Journal of Bacteriology, 189(22), 8034–8043. 10.1128/JB.00900-07

Guttenplan, S. B., & Kearns, D. B. (2013). Regulation of flagellar motility during biofilm formation. FEMS Microbiology Reviews, 37(6), 849–871. 10.1111/1574-6976.12018

Hirasawa, T., & Maeda, T. (2022). Adaptive Laboratory Evolution of Microorganisms: Methodology and Application for Bioproduction. Microorganisms, 11(1), 92. 10.3390/microorganisms11010092

Hufnagel, D. A., Depas, W. H., & Chapman, M. R. (2015). The Biology of the *Escherichia coli* Extracellular Matrix. Microbiology Spectrum, 3(3). 10.1128/microbiolspec.MB-0014-2014

Irizarry, R. A. (2003). Exploration, normalization, and summaries of high density oligonucleotide array probe level data. Biostatistics, 4(2), 249–264. 10.1093/biostatistics/4.2.249

Iyer, M. S., Pal, A., Srinivasan, S., Somvanshi, P. R., & Venkatesh, K. V. (2021). Global Transcriptional Regulators Fine-Tune the Translational and Metabolic Efficiency for Optimal Growth of Escherichia coli. MSystems, 6(2). 10.1128/msystems.00001-21

Kanehisa, M., Furumichi, M., Sato, Y., Kawashima, M., & Ishiguro-Watanabe, M. (2023). KEGG for taxonomy-based analysis of pathways and genomes. Nucleic Acids Research, 51(D1), D587–D592. 10.1093/nar/gkac963

Kavvas, E. S., Long, C. P., Sastry, A., Poudel, S., Antoniewicz, M. R., Ding, Y., Mohamed, E. T., Szubin, R., Monk, J. M., Feist, A. M., & Palsson, B. O. (2022). Experimental Evolution Reveals Unifying Systems-Level Adaptations but Diversity in Driving Genotypes. MSystems, 7(6). 10.1128/msystems.00165-22

Kawecki, T. J., Lenski, R. E., Ebert, D., Hollis, B., Olivieri, I., & Whitlock, M. C. (2012). Experimental evolution. Trends in Ecology & Evolution, 27(10), 547–560. 10.1016/j.tree.2012.06.001

Kim, K., Choe, D., Kang, M., Cho, S.-H., Cho, S., Jeong, K. J., Palsson, B., & Cho, B.-K. (2024). Serial adaptive laboratory evolution enhances mixed carbon metabolic capacity of Escherichia coli. Metabolic Engineering, 83, 160–171. 10.1016/j.ymben.2024.04.004

Kinnersley, M. A., Holben, W. E., & Rosenzweig, F. (2009). E Unibus Plurum: genomic analysis of an experimentally evolved polymorphism in Escherichia coli. PLoS Genetics, 5(11), e1000713. 10.1371/journal.pgen.1000713

Klemm, P., & Schembri, M. (2004). Type 1 Fimbriae, Curli, and Antigen 43: Adhesion, Colonization, and Biofilm Formation. EcoSal Plus, 1(1). 10.1128/ecosalplus.8.3.2.6

Kolde, R., Laur, S., Adler, P., & Vilo, J. (2012). Robust rank aggregation for gene list integration and meta-analysis. Bioinformatics, 28(4), 573–580. 10.1093/bioinformatics/btr709

Kram, K. E., Geiger, C., Ismail, W. M., Lee, H., Tang, H., Foster, P. L., & Finkel, S. E. (2017). Adaptation of Escherichia coli to Long-Term Serial Passage in Complex Medium: Evidence of Parallel Evolution. MSystems, 2(2). 10.1128/mSystems.00192-16

Kurokawa, M., & Ying, B.-W. (2019). Experimental Challenges for Reduced Genomes: The Cell Model Escherichia coli. Microorganisms, 8(1), 3. 10.3390/microorganisms8010003

Law, C. W., Chen, Y., Shi, W., & Smyth, G. K. (2014). voom: precision weights unlock linear model analysis tools for RNA-seq read counts. Genome Biology, 15(2), R29. 10.1186/gb-2014-15-2-r29

Lenski, R. E. (2017). Experimental evolution and the dynamics of adaptation and genome evolution in microbial populations. The ISME Journal, 11(10), 2181–2194. 10.1038/ismej.2017.69

Li, T., Liu, H., Lei, Q., & You, Z. (2026). Integrative transcriptomic meta-analysis reveals conserved transcriptional signatures and predictive biomarkers for active tuberculosis: a pathway-based machine learning approach. Frontiers in Microbiology, 17. 10.3389/fmicb.2026.1757941

Lisevich, I., Colin, R., Yang, H. Y., Ni, B., & Sourjik, V. (2025). Physics of swimming and its fitness cost determine strategies of bacterial investment in flagellar motility. Nature Communications, 16(1), 1731. 10.1038/s41467-025-56980-x

Love, M. I., Huber, W., & Anders, S. (2014). Moderated estimation of fold change and dispersion for RNA-seq data with DESeq2. Genome Biology, 15(12), 550. 10.1186/s13059-014-0550-8

McCloskey, D., Xu, S., Sandberg, T. E., Brunk, E., Hefner, Y., Szubin, R., Feist, A. M., & Palsson, B. O. (2018). Evolution of gene knockout strains of E. coli reveal regulatory architectures governed by metabolism. Nature Communications, 9(1), 3796. 10.1038/s41467-018-06219-9

Ni, B., Colin, R., Link, H., Endres, R. G., & Sourjik, V. (2020). Growth-rate dependent resource investment in bacterial motile behavior quantitatively follows potential benefit of chemotaxis. Proceedings of the National Academy of Sciences, 117(1), 595–601. 10.1073/pnas.1910849117

Ogasawara, H., Yamamoto, K., & Ishihama, A. (2011). Role of the Biofilm Master Regulator CsgD in Cross-Regulation between Biofilm Formation and Flagellar Synthesis. Journal of Bacteriology, 193(10), 2587–2597. 10.1128/JB.01468-10

Okano, H., Hermsen, R., Kochanowski, K., & Hwa, T. (2019). Regulation underlying hierarchical and simultaneous utilization of carbon substrates by flux sensors in Escherichia coli. Nature Microbiology, 5(1), 206–215. 10.1038/s41564-019-0610-7

Olivas-Bernal, C. A., Vargas-Albores, F., Garibay-Valdez, E., Cicala, F., & Martínez-Porchas, M. (2026). Transcriptomic Meta-Analysis as a Framework for Robust Cross-Study Biological Inference. International Journal of Molecular Sciences, 27(11), 4674. 10.3390/ijms27114674

Özel, A., Topaloğlu, A., Esen, Ö., Holyavkin, C., Baysan, M., & Çakar, Z. P. (2024). Transcriptomic and Physiological Meta-Analysis of Multiple Stress-Resistant Saccharomyces cerevisiae Strains. Stresses, 4(4), 714–733. 10.3390/stresses4040046

Patil, A. V., Zhao, J., Khairnar, S. V., Srivastava, S., Yang, L., & Anand, A. (2026). Cost adjusted hierarchical defense strategies enables stratified oxidative stress tolerance. 10.64898/2026.03.03.709455

Peng, W., Zhang, X., Qi, Q., & Liang, Q. (2025). Advances in adaptive laboratory evolution applications for Escherichia coli. Synthetic and Systems Biotechnology, 10(4), 1306–1321. 10.1016/j.synbio.2025.07.011

Pesavento, C., Becker, G., Sommerfeldt, N., Possling, A., Tschowri, N., Mehlis, A., & Hengge, R. (2008). Inverse regulatory coordination of motility and curli-mediated adhesion in *Escherichia coli*. Genes & Development, 22(17), 2434–2446. 10.1101/gad.475808

Prigent-Combaret, C., Brombacher, E., Vidal, O., Ambert, A., Lejeune, P., Landini, P., & Dorel, C. (2001). Complex Regulatory Network Controls Initial Adhesion and Biofilm Formation in *Escherichia coli* via Regulation of the *csgD* Gene. Journal of Bacteriology, 183(24), 7213–7223. 10.1128/JB.183.24.7213-7223.2001

Puentes-Téllez, P. E., Kovács, Á. T., Kuipers, O. P., & van Elsas, J. D. (2014). Comparative genomics and transcriptomics analysis of experimentally evolved Escherichia coli MC1000 in complex environments. Environmental Microbiology, 16(3), 856–870. 10.1111/1462-2920.12239

Ritchie, M. E., Phipson, B., Wu, D., Hu, Y., Law, C. W., Shi, W., & Smyth, G. K. (2015). limma powers differential expression analyses for RNA-sequencing and microarray studies. Nucleic Acids Research, 43(7), e47–e47. 10.1093/nar/gkv007

Rossi, E., Paroni, M., & Landini, P. (2018). Biofilm and motility in response to environmental and host-related signals in Gram negative opportunistic pathogens. Journal of Applied Microbiology, 125(6), 1587–1602. 10.1111/jam.14089

Rychel, K., Chen, K., Catoiu, E. A., Olson, E., Sandberg, T. E., Gao, Y., Xu, S., Hefner, Y., Szubin, R., Patel, A., Feist, A. M., & Palsson, B. O. (2025). Laboratory Evolution Reveals Transcriptional Mechanisms Underlying Thermal Adaptation of *Escherichia coli*. Genome Biology and Evolution, 17(10). 10.1093/gbe/evaf171

Sandberg, T. E., Salazar, M. J., Weng, L. L., Palsson, B. O., & Feist, A. M. (2019). The emergence of adaptive laboratory evolution as an efficient tool for biological discovery and industrial biotechnology. Metabolic Engineering, 56, 1–16. 10.1016/j.ymben.2019.08.004

Saxena, P., Samanta, D., Thakur, P., Gopalakrishnan, V., & Sani, R. K. (2025). Comparative transcriptomics analysis of the Oleidesulfovibrio alaskensis G20 biofilms grown on copper and polycarbonate surfaces. Biofilm, 10, 100309. 10.1016/j.bioflm.2025.100309

Schwartz, K., Kinnersley, M., Lindsey, C. R., Sherlock, G., & Rosenzweig, F. (2025). Adaptive genetics reveals constraints on protein structure/function by evolving E. coli under constant nutrient limitation. BMC Biology, 23(1), 261. 10.1186/s12915-025-02331-7

Serra, D. O., & Hengge, R. (2021). Bacterial Multicellularity: The Biology of *Escherichia coli* Building Large-Scale Biofilm Communities. Annual Review of Microbiology, 75(1), 269–290. 10.1146/annurev-micro-031921-055801

Shannon, P., Markiel, A., Ozier, O., Baliga, N. S., Wang, J. T., Ramage, D., Amin, N., Schwikowski, B., & Ideker, T. (2003). Cytoscape: A Software Environment for Integrated Models of Biomolecular Interaction Networks. Genome Research, 13(11), 2498–2504. 10.1101/gr.1239303

Szklarczyk, D., Kirsch, R., Koutrouli, M., Nastou, K., Mehryary, F., Hachilif, R., Gable, A. L., Fang, T., Doncheva, N. T., Pyysalo, S., Bork, P., Jensen, L. J., & von Mering, C. (2023). The STRING database in 2023: protein–protein association networks and functional enrichment analyses for any sequenced genome of interest. Nucleic Acids Research, 51(D1), D638–D646. 10.1093/nar/gkac1000

Thomason, M. K., Fontaine, F., De Lay, N., & Storz, G. (2012). A small RNA that regulates motility and biofilm formation in response to changes in nutrient availability in *Escherichia coli*. Molecular Microbiology, 84(1), 17–35. 10.1111/j.1365-2958.2012.07965.x

Travisano, M., Mongold, J. A., Bennett, A. F., & Lenski, R. E. (1995). Experimental Tests of the Roles of Adaptation, Chance, and History in Evolution. Science, 267(5194), 87–90. 10.1126/science.7809610

Tursi, S. A., & Tükel, Ç. (2018). Curli-Containing Enteric Biofilms Inside and Out: Matrix Composition, Immune Recognition, and Disease Implications. Microbiology and Molecular Biology Reviews, 82(4). 10.1128/MMBR.00028-18

van Heeswijk, W. C., Westerhoff, H. V., & Boogerd, F. C. (2013). Nitrogen Assimilation in Escherichia coli: Putting Molecular Data into a Systems Perspective. Microbiology and Molecular Biology Reviews, 77(4), 628–695. 10.1128/MMBR.00025-13

Wang, G., Li, Q., Zhang, Z., Yin, X., Wang, B., & Yang, X. (2023). Recent progress in adaptive laboratory evolution of industrial microorganisms. Journal of Industrial Microbiology and Biotechnology, 50(1). 10.1093/jimb/kuac023

Wang, X., Preston, J. F., & Romeo, T. (2004). The *pgaABCD* Locus of *Escherichia coli* Promotes the Synthesis of a Polysaccharide Adhesin Required for Biofilm Formation. Journal of Bacteriology, 186(9), 2724–2734. 10.1128/JB.186.9.2724-2734.2004

Wang, X., Zorraquino, V., Kim, M., Tsoukalas, A., & Tagkopoulos, I. (2018). Predicting the evolution of Escherichia coli by a data-driven approach. Nature Communications, 9(1), 3562. 10.1038/s41467-018-05807-z

Wu, T., Hu, E., Xu, S., Chen, M., Guo, P., Dai, Z., Feng, T., Zhou, L., Tang, W., Zhan, L., Fu, X., Liu, S., Bo, X., & Yu, G. (2021). clusterProfiler 4.0: A universal enrichment tool for interpreting omics data. The Innovation, 2(3), 100141. 10.1016/j.xinn.2021.100141

Zaykin, D. V. (2011). Optimally weighted Z-test is a powerful method for combining probabilities in meta-analysis. Journal of Evolutionary Biology, 24(8), 1836–1841. 10.1111/j.1420-9101.2011.02297.x

Zhong, S., Miller, S. P., Dykhuizen, D. E., & Dean, A. M. (2009). Transcription, Translation, and the Evolution of Specialists and Generalists. Molecular Biology and Evolution, 26(12), 2661–2678. 10.1093/molbev/msp187

Zion, S., Katz, S., & Hershberg, R. (2024). Escherichia coli adaptation under prolonged resource exhaustion is characterized by extreme parallelism and frequent historical contingency. PLOS Genetics, 20(6), e1011333. 10.1371/journal.pgen.1011333

